# An In Vitro BRAF Activation Assay Elucidates Molecular Mechanisms Driving Disassembly of the Autoinhibited BRAF State

**DOI:** 10.1101/2025.08.19.671159

**Authors:** Daniel A. Ritt, David E. Durrant, Matthew R. Drew, Suzanne I. Sandin, Juliana A. Martinez Fiesco, Xiaohua Zhang, Fikret Aydin, Timothy S. Carpenter, Grace M. Scheidemantle, Robert A. D’Ippolito, Alexandria L. Sohn, Caroline J. DeHart, Rebika Shrestha, Kelly Snead, Jenna Hull, Jeremy O. B. Tempkin, Yue Yang, Felice C. Lightstone, Frederick H. Streitz, Ping Zhang, Thomas J. Turbyville, Andrew G. Stephen, Dominic Esposito, Helgi I. Ingolfsson, Dwight V. Nissley, Deborah K. Morrison

## Abstract

The RAF kinases (ARAF, BRAF and CRAF) are essential components of the RAS-ERK signaling pathway, which controls vital cellular processes and is frequently dysregulated in human disease. Notably, mutations that alter BRAF function are prominent drivers of human cancer and certain RASopathy disorders, making BRAF an important target for therapeutic intervention. Despite extensive research, several aspects of BRAF regulation remain unclear. In this study, we developed an in vitro BRAF activation assay using purified autoinhibited BRAF:14-3-3_2_:MEK complexes. Our results show that fully processed, active-state KRAS alone can promote dimer-dependent BRAF activation. Moreover, we found that phosphatidylserine (PS)-containing liposomes synergized with KRAS to promote BRAF activation, achieving activity levels comparable to those observed with BRAF proteins that constitutively dimerize. In contrast, the SMP phosphatase complex had only a minimal effect on BRAF catalytic activity in this system but mediated the dephosphorylation of the negative regulatory pS365 14-3-3 binding site in a manner that was accelerated by the presence of KRAS alone or KRAS and 30% PS liposomes. Finally, we show that inhibitors blocking the BRAF RBD:KRAS interaction were able to suppress the in vitro activation of BRAF, underscoring the critical role of RAS binding in initiating the disassembly of the BRAF autoinhibited state. Thus, this assay provides valuable insights into the steps required for BRAF activation and can serve as an effective screening tool for identifying compounds that may inhibit this process and have therapeutic potential.

**Significance Statement:** BRAF is a central intermediate in RAS pathway signaling, and its activity is often elevated in human cancers and RASopathy disorders. Due to the complexity of BRAF activation, identifying compounds that sustainably inhibit BRAF function has proven difficult, emphasizing the need for a more comprehensive understanding of BRAF regulation. Here, we have developed an in vitro BRAF activation assay that elucidates key steps in this process. Our findings demonstrate that RAS binding not only recruits BRAF to the plasma membrane but initiates the disassembly of the autoinhibited monomer, which in the context of the membrane, facilitates BRAF dimerization and activation. This assay advances our understanding of BRAF regulation and provides a novel platform for drug discovery efforts targeting BRAF.

## Introduction

The RAF kinases—ARAF, BRAF, and CRAF—play a central role in cell signaling, acting as direct effectors of the RAS GTPases and as the initiating kinase in the ERK cascade, comprised of the RAF, MEK, and ERK protein kinases (1). The RAFs function in the transmission of signals required for normal growth and development as well as those contributing to human disease, including cancer and the RASopathy developmental disorders (2–4).

The RAF kinases can be divided into two functional domains: the C-terminal kinase domain and the N-terminal regulatory domain (1, 5). The regulatory domain contains a variable-length N-terminal segment, followed by the <u>R</u>AS-<u>b</u>inding <u>d</u>omain (RBD), a <u>c</u>ysteine-<u>r</u>ich, zinc finger <u>d</u>omain (CRD), and a linker region. The RBD serves as the high-affinity binding site for RAS (6), whereas the CRD interacts with the plasma membrane and RAS during RAF activation but helps maintain RAF in an inactive state under quiescent conditions (7–13). The RAF kinases also possess two conserved, phosphorylation-dependent binding sites for 14-3-3 dimers: one in the linker region of the regulatory domain (N’ site) and another in the C-terminal tail (C’ site) following the kinase domain (14).

In resting cells, the RAFs localize to the cytosol as inactive monomers (15). For the RAFs to become active enzymes, dimerization of the kinase domains is required under most signaling conditions (16). Studies have further shown that the RAF kinases selectively interact with GTP-bound RAS (17–19), and that contact with RAS and the plasma membrane is typically required for RAF dimerization and activation (10, 20, 21). More recently, valuable insights regarding RAF activation have emerged from cryo-electron microscopy (cryo-EM) structures of full-length BRAF complexes. In particular, structures of the monomeric BRAF:14-3-3_2_:MEK complex show that, in the inactive monomeric state, a 14-3-3 dimer can bind simultaneously to both the N’ and C’ sites, obscuring the membrane-binding loops of the CRD and the kinase domain “RKTR” motif, which is critical for dimerization (22–24). These structures also underscore the central role of the CRD in maintaining BRAF in the autoinhibited state, through interactions with the kinase domain and both 14-3-3 protomers. In contrast, domains in the regulatory region were not resolved in the structures of active dimers (24, 25), likely due to the flexibility of this region in the absence of membrane-binding partners. However, the kinase domains of the BRAF dimer were clearly resolved, with a single 14-3-3 dimer bridging the two kinase domains at the C’ site. Collectively, these structures indicate that substantial movement of the BRAF domains, coupled with a rearrangement in 14-3-3 binding is required for BRAF to transition from an autoinhibited monomer to an active dimer.

Although studies have confirmed that RAS binding, interactions with the plasma membrane, and dephosphorylation of the N’ 14-3-3 binding site by the SHOC2/MRAS/PP1 (SMP) complex occur during the RAF activation process (5), the precise molecular mechanisms by which these events modulate RAF are not fully understood. For example, while high-affinity binding to membrane-bound RAS is mediated by the RAF RBD, it is unclear whether the RBD:RAS interaction functions solely to recruit BRAF to the plasma membrane or whether it actively contributes to the disruption of the autoinhibited state. In this study, we have developed an in vitro BRAF activation assay utilizing purified autoinhibited BRAF:14-3-3_2_:MEK complexes. This assay has allowed us to investigate the role of key components implicated in the BRAF monomer-to-dimer transition.

## Results

### Analysis of the BRAF RBD:14-3-3 Interface

Previously, we reported cryo-EM structures of two autoinhibited complexes, BRAF:14-3-3_2_:MEK and BRAF:14-3-3_2_, both of which were isolated from serum-depleted human 293FT cells (24). In these structures, the RBD was well-resolved and positioned atop the 14-3-3 protomer bound to the C^′^ pS729 site, generating an extensive contact surface (~435 Å^2^) with considerable charge complementarity. To further investigate the significance of the RBD:14-3-3 interface, all-atom molecular dynamics simulations were performed using the autoinhibited BRAF structure (PDB ID:7MFD). In 135 independent simulations, with a cumulative time of 73.2 microseconds, the RBD consistently maintained contact with the 14-3-3 protomer (SI appendix, Fig. S1*A* and Movie S1). Numerous hydrogen bonds were observed between residues in the α1-helix and loop 3 of the RBD and residues in the α8-helix, α9-helix, and loop 8 of 14-3-3 (Fig. 1*A*). Notably, the RBD residue K182 formed hydrogen bonds with several 14-3-3 residues, including E208, L206, and Y211, and was involved in bond formation 68% of the simulation time. These findings suggest that bond formation at the RBD:14-3-3 interface may contribute to the stability of the autoinhibited conformation and help orient the RBD for RAS binding.

**Fig 1.**
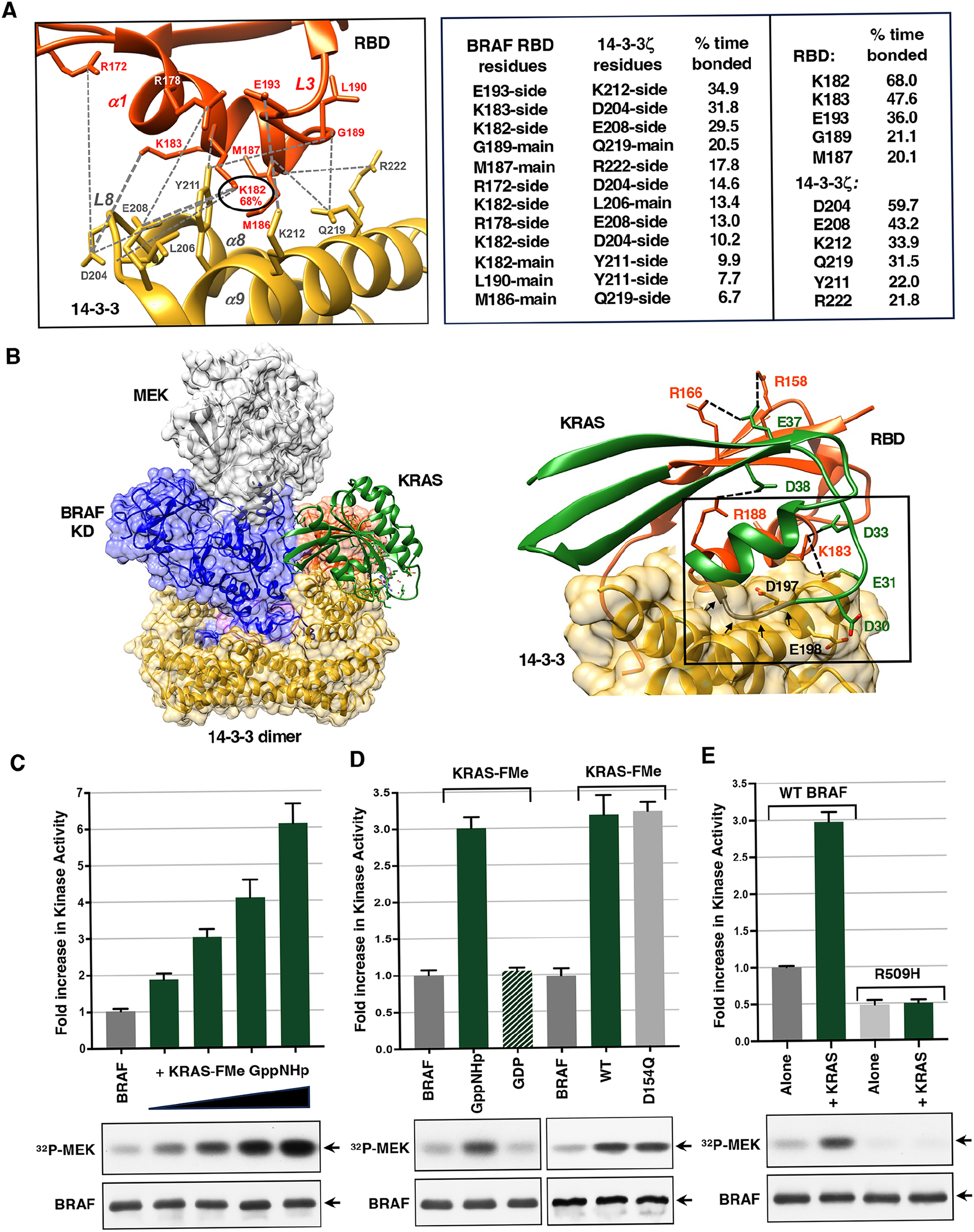
KRAS binding disrupts the BRAF RBD:14-3-3 interface and promotes BRAF activation in vitro. *(A)* Analysis of the BRAF RBD:14-3-3 interface is shown (left), highlighting residues of the RBD (red) and 14-3-3 (yellow) that form hydrogen bonds (dashed gray lines) during molecular dynamics simulations of the BRAF autoinhibited complex (PBD ID: 7MFD). The RBD K182 residue (circled) exhibited bond formation the highest percentage of simulation time. The percentage of time (greater than 5%) that bond formation was observed between specific RBD and 14-3-3 residues is shown in the center, while the percentage of time (greater than 20%) that individual RBD or 14-3-3 residues were bonded is presented on the right. *(B)* Cryo-EM structure of the autoinhibited BRAF:14-3-3_2_:MEK complex (PBD ID: 7MFD) is shown with the BRAF RBD in red, the BRAF kinase domain (KD) in blue, 14-3-3 in yellow, and MEK in gray. KRAS binding is modeled onto the autoinhibited BRAF structure (left). Formation of critical ionic bonds (indicated by dashed black lines) between the RBD (R158, R166, K183, and R188) and KRAS (D38, E37, D33, and E31) residues is predicted to cause electrostatic repulsion between acidic residues of KRAS (D30 and E31) and 14-3-3 (E198 and D197) and steric clash (indicated with black arrows) between the KRAS switch I loop and the α8 and α9 helices of 14-3-3 (right). (*C)* Autoinhibited BRAF complexes were incubated alone or in the presence of increasing amounts of GppNHp-bound KRAS-FMe (5, 10, 15, and 20 molar excess in comparison to BRAF) for 1 hr and then monitored for kinase activity in vitro using kinase-dead MEK as a substrate. *(D)* Autoinhibited BRAF complexes were incubated alone or in the presence of GppNHp- or GDP-bound WT KRAS-FMe or GppNHp-bound D154Q KRAS-FMe (10 molar excess) prior to monitoring for kinase activity. *(E)* Autoinhibited WT or dimerization-defective R509H BRAF complexes were incubated in the presence or absence of GppNHp-KRAS-FMe (10 molar excess) prior to monitoring for kinase activity. *(C-E)* The graphs indicate the average fold increase in kinase activity, with BRAF activity alone set at 1, based on data from at least 3 independent experiments ± SD. Also shown are autoradiographs of the ^32^P-labeled MEK and immunoblot analyses of BRAF levels from representative experiments.

Indeed, in autoinhibited BRAF structures from mammalian cells, the RBD is positioned such that the four basic residues—R158, R166, K183, and R188—that form critical ionic bonds with the switch I region of KRAS are either fully exposed (R158, R166, R188) or partially exposed (K183) (SI appendix, Fig. S1*B*). However, superimposing KRAS onto the autoinhibited BRAF structure indicates that forming bonds with these residues would result in steric clashes between the KRAS switch I loop and the α8- and α9-helices of 14-3-3 and would generate electrostatic repulsion between acidic residues in KRAS, D30 and E31, and 14-3-3, D197 and E198 (Fig. 1*B*). Based on these observations, it has been proposed that RBD:RAS binding may initiate the disassembly of the autoinhibited state (24).

### In Vitro Activation of Autoinhibited BRAF Complexes by KRAS-FMe

To test the above hypothesis and to assess the contribution of other components implicated in RAF activation, we developed an in vitro BRAF activation assay using purified components (SI appendix, Fig. S2*A*). For this assay, we utilized autoinhibited BRAF:14-3-3_2_:MEK complexes isolated from 293FT cells that were highly phosphorylated on the pS365 and pS729 14-3-3 binding sites (97.83% for pS365 and 99.57% for pS729 SI appendix Fig. S2*B* and Dataset S1). The BRAF complexes were incubated in vitro for one hour with or without purified full-length KRAS, which was farnesylated and methylated at the C-terminus (KRAS-FMe), and bound to the non-hydrolyzable GTP analog, GppNHp. For some experiments, phosphatidylserine (PS)-containing liposomes and/or the SMP phosphatase complex were added to the BRAF complexes in the presence or absence of KRAS. Following the incubation period, the samples were monitored for changes in BRAF activity using in vitro kinase assays with kinase-dead MEK as a substrate. It is important to note that the basal activity of the autoinhibited BRAF complexes alone remained unchanged throughout the one-hour incubation reaction (Fig. S2*C*), indicating that the observed changes in BRAF activity can be directly attributed to the addition of components to the incubation reaction.

As shown in Fig. 1*C*, the kinase activity of BRAF increased in a dose-dependent manner when incubated with increasing amounts of GppNHp-bound KRAS-FMe. Activation of the BRAF complexes required that KRAS be in the active state, as incubation with GDP-bound KRAS-FMe resulted in little to no change in BRAF activity (Fig. 1*D*). Moreover, the in vitro activation of BRAF appeared to be driven by RAS-induced BRAF dimerization, as autoinhibited BRAF complexes containing the dimer-disrupting R509H mutation exhibited lower basal activity and showed no activation upon incubation with GppNHp-KRAS-FMe (Fig. 1*E*). In contrast, GppNHp-KRAS-FMe containing the D154Q mutation—proposed to impair RAS dimerization (Ambrogio et al., 2018)—was fully competent to activate BRAF (Fig. 1*D*), consistent with a model in which RAS proteins function through membrane clustering rather than discrete dimerization. These findings suggest that binding to active-state KRAS-FMe promotes disassembly of the autoinhibited BRAF state and facilitates the formation of active BRAF dimers.

### Effect of Liposomes on BRAF Activation In Vitro

During RAS-mediated signaling, the RAF CRD interacts with RAS and binds to negatively charged lipids, particularly phosphatidylserine (PS), in the plasma membrane (8-10, 12). Moreover, interactions between the CRD and the plasma membrane are required for full RAF activation (10, 21). Therefore, to further investigate the role of the plasma membrane in RAF activation, we next examined whether incubation with PS-containing liposomes would alter BRAF activity, either alone or in the presence of GppNHp-KRAS-FMe (hereafter referred to as KRAS). As shown in Fig. 2*A*, the addition of liposomes containing 30% PS resulted in a modest, dose-dependent increase in BRAF activity; however, the level of activation was markedly enhanced if KRAS was also present. When we examined the effect of varying the PS composition of the liposomes, we found that liposomes containing 0% or 15% PS had little effect on BRAF activity, whether added alone or in combination with KRAS (Fig. 2*B*). In contrast, BRAF activity was elevated when incubated with liposomes containing 30% or 50% PS, and both types of liposomes synergized with KRAS to promote BRAF activation (Fig. 2*B*). Finally, when we investigated whether 8-lipid liposomes, which are more membrane-like (26, 27), could further enhance KRAS-mediated BRAF activation in the in vitro assay, we observed no significant difference in the activation levels achieved whether 8-lipid or 30% PS liposomes were added in combination with KRAS (Fig. 2*C*).

**Fig. 2.**
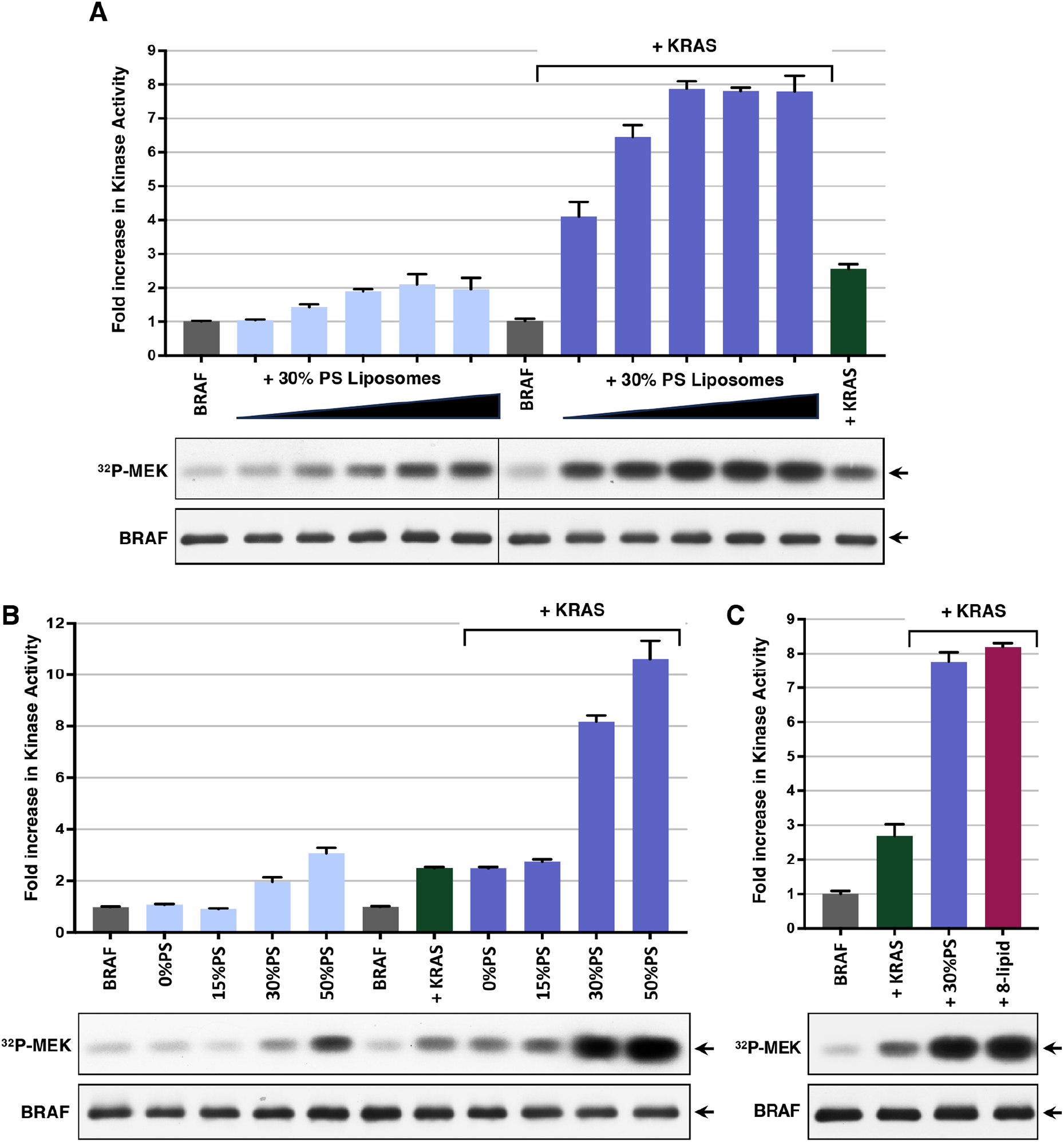
PS-containing liposomes synergize with KRAS to promote BRAF activation in vitro. *(A)* Autoinhibited BRAF complexes were incubated alone or with increasing amounts of 30% PS-containing liposomes (20.5-328 μM) in the presence or absence of GppNHp-KRAS-FMe (7 molar excess compared to BRAF) prior to monitoring for kinase activity. *(B)* Kinase activity of autoinhibited BRAF complexes incubated alone or with liposomes containing 0, 15, 30 or 50% PS (41 μM) in the presence or absence of GppNHp-KRAS-FMe (7 molar excess) is shown. *(C)* Kinase activity of autoinhibited BRAF complexes incubated alone or with 30% PS or 8-lipid liposomes (41 μM) in the presence of GppNHp-KRAS-FMe (7 molar excess). *(A-C)* As a control, the kinase activity of autoinhibited BRAF complexes incubated with GppNHp-KRAS-FMe alone is included. The graphs represent the average fold increase in kinase activity, with BRAF activity alone set at 1, based on data from 3 independent experiments ± SD. Also shown are autoradiographs of the ^32^P-labeled MEK and immunoblot analyses of BRAF levels from representative experiments.

### Effect of the SMP Phosphatase Complex on BRAF Activation In Vitro

SMP-mediated dephosphorylation of the N^′^ 14-3-3 binding site (pS365 for BRAF, pS259 for CRAF, and pS214 for ARAF) occurs when RAF is recruited to the plasma membrane for activation (28–31). In addition, CRAF mutations disrupting the phosphorylation of this site upregulate the biological activity of CRAF and are disease drivers in the RASopathy disorder Noonan Syndrome (32, 33). Therefore, to determine whether the SMP complex and dephosphorylation of the BRAF pS365 site can directly modulate BRAF’s catalytic activity, experiments were performed in which soluble SMP complexes were added to the autoinhibited BRAF complexes, either alone or in combination with KRAS, 30% liposomes, or both KRAS and 30% liposomes. As shown in Fig. 3*A*, the addition of SMP to the incubation reaction caused only minimal increases (:Σ10%) in the activity levels induced by KRAS and/or 30% PS liposomes. However, dephosphorylation of the BRAF pS365 N^′^ site was detected and only observed when SMP was present in the incubation reactions (Fig. *3A* and SI appendix, Fig. 3*A-B*). Of note, a more pronounced activating effect of SMP was observed when suboptimal amounts of KRAS (3 molar excess) are used in the activation assay (~40% increase; SI appendix, Fig. S3*B*), suggesting that SMP function may be more critical under conditions of low KRAS activation.

**Fig. 3.**
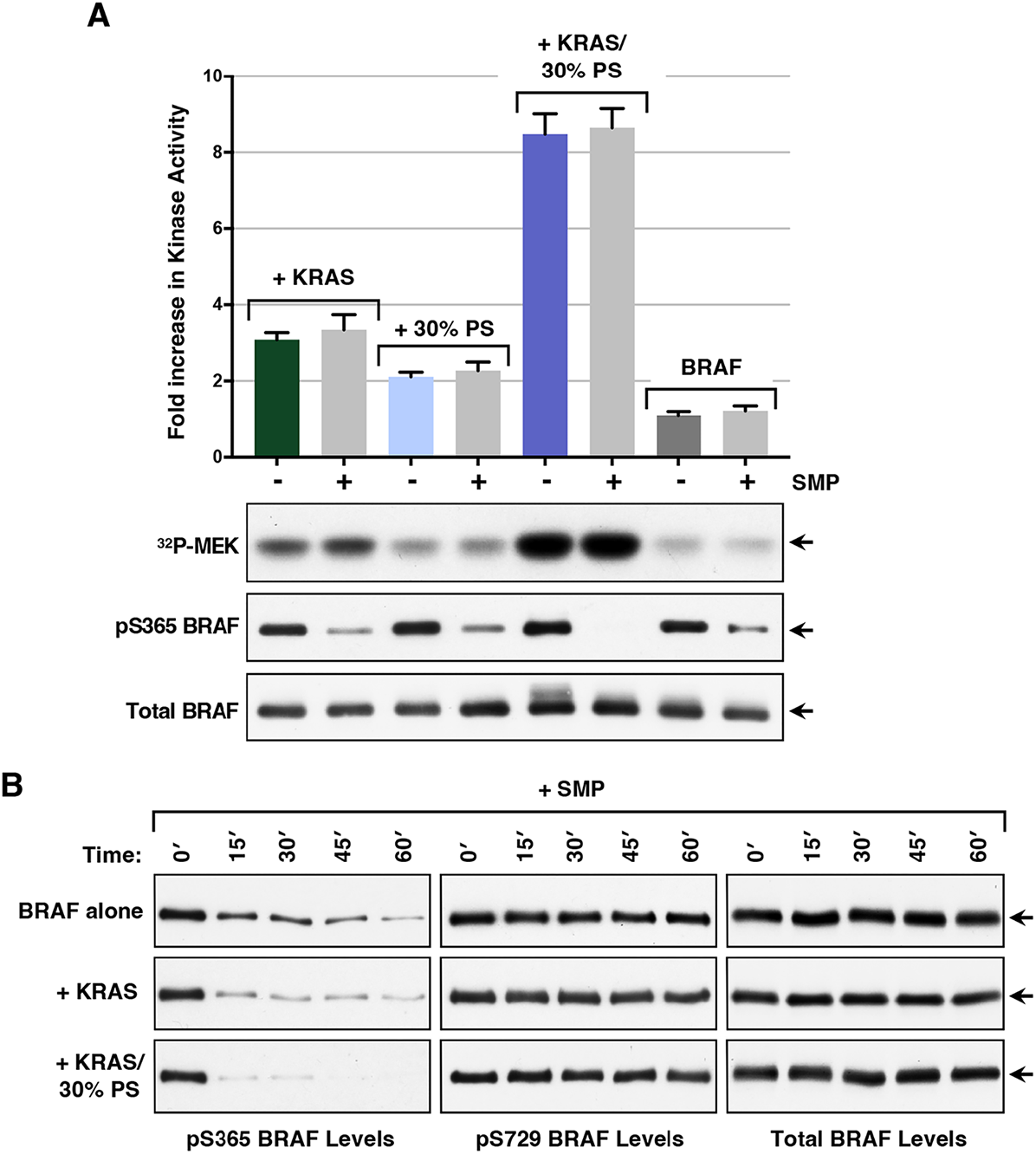
Effect of SMP in the in vitro BRAF activation assay. *(A)* Autoinhibited BRAF complexes were incubated alone or with the indicated components prior to monitoring for kinase activity. The graph represents the average fold increase in kinase activity, with BRAF activity alone set at 1, based on data from at least 3 independent experiments ± SD. Also shown are autoradiographs of the ^32^P-labeled MEK and immunoblot analyses of pS365 levels and total BRAF levels from a representative experiment. *(B)* Autoinhibited BRAF complexes were incubated with SMP alone for the indicated times or were incubated with SMP in the presence of GppNHp-KRAS- or GppNHp-KRAS-FMe and 30% PS liposomes Samples were then examined by immunoblot analysis for pS365 BRAF levels and total BRAF levels. For these assays, KRAS-FMe was used at a 7 molar excess with respect to BRAF, SMP at 1.2 molar excess, and 30% PS liposomes at a concentration of 83 μM.

Because dephosphorylation of the pS365 site appeared more complete when KRAS and 30% PS liposomes were present, we next examined the dephosphorylation of this site as a function of time during the incubation period (Fig. 3*B*). Our findings indicate that dephosphorylation of this site occurred in a time-dependent manner when SMP was added alone to the BRAF complexes, suggesting a dynamic association between the 14-3-3 protomer and this site. Strikingly, dephosphorylation of the pS365 site was more efficient when KRAS was present and was further accelerated in the presence of both KRAS and 30% liposomes. These findings support the model that KRAS binding, augmented by contact with membrane lipids, can promote the disruption of the autoinhibited BRAF complex, thereby increasing the exposure of the pS365 site for dephosphorylation.

### Analysis of BRAF Monomer Complexes Produced in Sf9 and Tni-FNL Insect Cells

The RBD was well-resolved in cryo-EM structures of autoinhibited BRAF:14-3-3_2_:MEK complexes isolated from serum-depleted human 293FT cells (24). However, in structures of BRAF:14-3-3_2_:MEK complexes produced in Sf9 (*Spodoptera frugiperda*) insect cells, the RBD was not clearly resolved, and in subsequent cryo-EM structures of Sf9-produced complexes bound to the KRAS G-domain, no contact was observed between the RBD and 14-3-3, suggesting an increased mobility of the RBD (23, 34). Additionally, soluble KRAS or lipid nanodiscs containing KRAS were reported to have only a modest effect on the activity of Sf9-produced BRAF:14-3-3_2_:MEK complexes in in vitro kinase assays (34, 35). To determine whether similar effects would be observed for Sf9-produced BRAF complexes generated using our production and purification protocols, human BRAF, MEK1, and 14-3-3ζ were co-expressed in Sf9 insect cells, and BRAF:14-3-3_2_:MEK complexes were isolated. For comparison, the BRAF:14-3-3_2_:MEK complexes were also produced in the Tni-FNL (*Trichoplusia ni*) insect cell system. Purified Sf9 and Tni-FNL-produced complexes (SI appendix, Fig. S4*A*) were then assessed for their enzymatic activity in the BRAF activation assay and for their phosphorylation state using mass spectrometry.

As shown in Fig. 4*A*, the basal activity of the Sf9 complexes alone was significantly higher than that of the Tni-FNL complexes. Further analysis revealed that the kinase activity of the Sf9-produced complexes alone increased over time during the incubation reaction, whereas the activity of the Tni-FNL-produced complexes remained constant throughout the incubation period (SI appendix, Fig. S4*B*), suggesting that the Tni-FNL complexes adopt a more stable autoinhibited conformation. When the BRAF complexes were incubated with KRAS and 30% PS liposomes, the Sf9-produced complexes exhibited only a modest 2-3 fold increase in activity (Fig. 4*B*), consistent with previous reports (34, 35). In contrast, the basal activity of the Tni-FNL-produced complexes was comparable to that of the 293FT-produced complexes and increased 8-10 fold, in a manner similar to that observed for the complexes isolated from serum-depleted human 293FT cells (Fig. 4*A-B* and SI appendix, Fig.S4*C*).

**Fig. 4.**
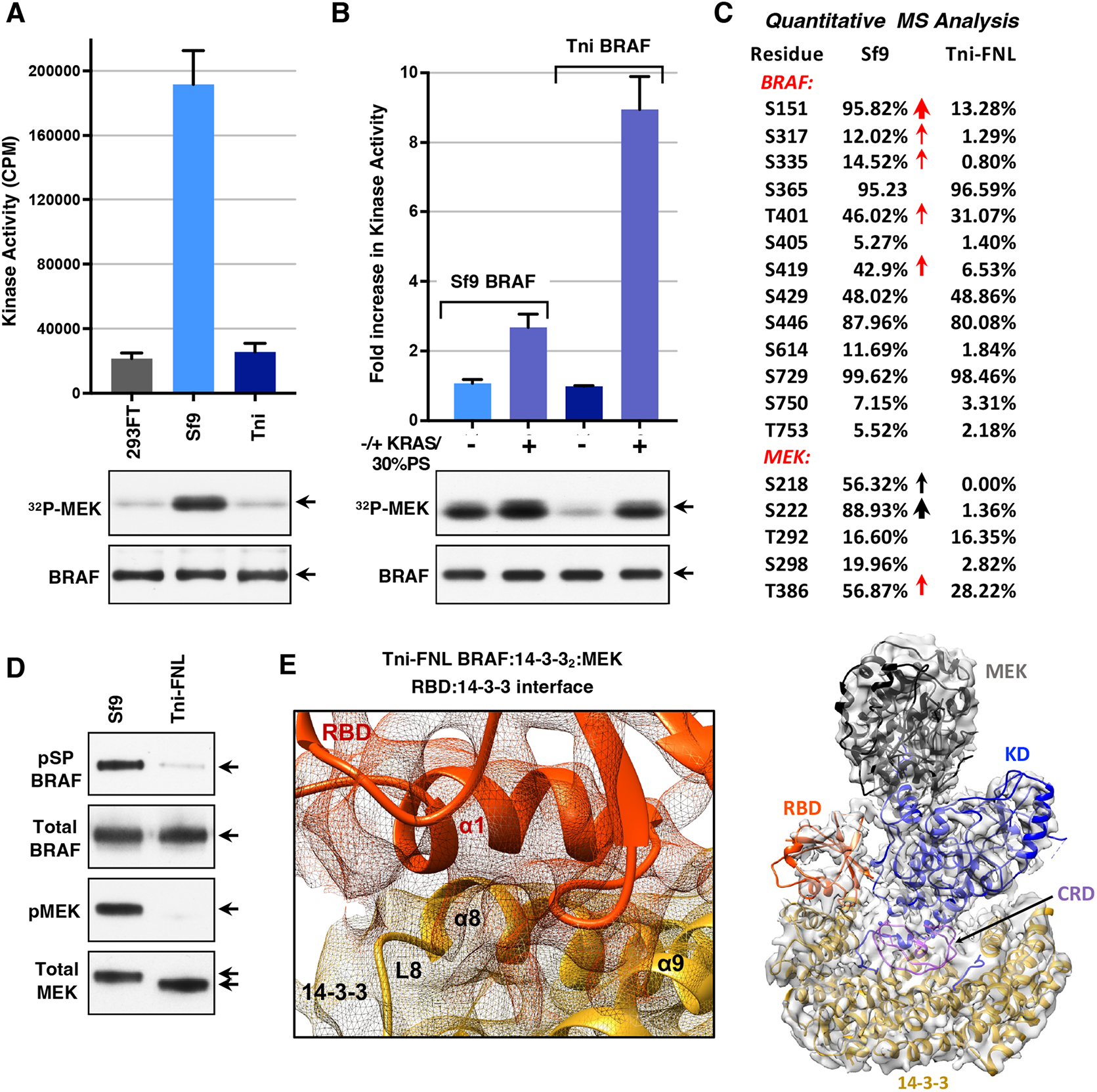
Analysis of Sf9- and Tni-FNL-produced BRAF:14-3-3_2_:MEK complexes. *(A)* BRAF:14-3-3_2_:MEK complexes produced in Sf9 and Tni-FNL insect cells or in human 293FT cells were monitored for kinase activity in vitro. The graph indicates the average CPM incorporated into kinase-dead MEK from 3 independent experiments ± SD. *(B)* The SF9 and Tni-FNL-produced BRAF complexes were incubated alone or with GppNHp-KRAS-FMe (7 molar excess) and 30% PS liposomes (83 μM) for 1 hr prior to monitoring for kinase activity. The graph represents the average fold increase in kinase activity, with BRAF activity alone set at 1, based on data from 3 independent experiments ± SD. Also shown are autoradiographs of the ^32^P-labeled MEK and immunoblot analyses of BRAF levels from representative experiments (*A-B*). *(C)* Sf9- and Tni-FNL-produced BRAF:14-3-3_2_:MEK complexes were evaluated for the phosphorylation state of BRAF and MEK using peptide reaction monitoring mass spectrometry analysis. Shown are residues where the relative percentage of phosphorylation was above 5%. Red arrows indicate pS/TP sites with >10% increased phosphorylation in the Sf9 versus Tni-FNL complexes. *(D)* Sf9- and Tni-FNL produced BRAF:14-3-3_2_:MEK complexes were examined by immunoblot analysis for pSP BRAF, total BRAF, pMEK (pS218/pS222), and total MEK levels. *(E)* Cryo-EM structure of Tni-FNL-produced BRAF:14-3-3_2_:MEK complexes, with the RBD forming a similar RBD:14-3-3 interface as observed in 293FT BRAF:14-3-3_2_:MEK structures. Same protein coloring as in Fig. 1*B*.

In addition to differences in activity, significant variations in the phosphorylation states of the Sf9 and Tni-FNL BRAF:14-3-3_2_:MEK complexes were also observed, with the BRAF and MEK proteins in the Sf9 complexes exhibiting higher levels of phosphorylation (Fig. 4*C-D* and SI appendix, Fig. S5, S6, and Dataset S2). In particular, phosphorylation of MEK1 at the activation segment S218/S222 sites was high in the Sf9 complexes, whereas little to no phosphorylation at these sites was detected in the Tni-FNL complexes (Fig. 4*C-D* and SI appendix, Dataset S2). Phosphorylation of both MEK1 and BRAF at known ERK-mediated sites (T386 for MEK1 and S151 and S419 for BRAF) was also elevated in the Sf9 complexes, with the levels of phosphorylated S151 BRAF reaching 95% in the Sf9 complexes, compared to only 13% in the Tni-FNL complexes (Fig. 4*C* and SI appendix, Dataset S2). The S151 site immediately precedes the RBD, and the high levels of phosphorylation at this residue may explain the increased RBD mobility indicated in cryo-EM structures of Sf9-produced BRAF:14-3-3_2_:MEK complexes (23, 34). Consistent with these findings, all-atom molecular dynamics simulations of the Sf9-produced BRAF:14-3-3_2_:MEK structure (638 simulations with an aggregate time of over 406 microseconds for PDB ID: 6NYB) also revealed a high degree of RBD mobility (SI appendix, Fig. S7*A*). However, spontaneous, unbiased association of the RBD with the 14-3-3 protomer bound to the pS729 site could be observed, with the RBD settling on the 14-3-3 protomer to form similar, but slightly distinct, contacts compared to those seen in the 293FT-produced autoinhibited BRAF complex (SI appendix, Fig. S7*B-C* and Movie S2).

Because the BRAF complexes derived from 293FT and Tni-FNL cells exhibited similar responses in the in vitro BRAF activation assays, the Tni-FNL-BRAF complexes were subjected to cryo-EM analysis to determine if their configuration was the same as that of the autoinhibited complexes from 293FT cells. This analysis yielded a structure of the Tni-FNL BRAF:14-3-3_2_:MEK complex at a resolution of 3.8 Å. When aligned with the 293FT BRAF:14-3-3_2_:MEK structure, the two complexes showed a Cα R.M.S.D. of 0.36 Å (Fig. 4*E* and SI appendix, Fig. S8*A-G*). The RBD was resolved and assumed the same position as observed in the 293FT BRAF:14-3-3_2_:MEK structure, with a similar interface between the RBD and the 14-3-3 protomer bound to the pS729 site (Fig. 4*E*). These findings indicate that the Tni-FNL BRAF:14-3-3_2_:MEK complexes accurately represent the full autoinhibited state observed for complexes isolated from serum-depleted human 293FT cells.

### Utilization of the In Vitro BRAF Activation Assay

Given that the Tni-FNL expression system allows for increased production of fully autoinhibited BRAF complexes, we used this system to further evaluate the utility of the in vitro BRAF activation assay. To determine whether the level of BRAF activity achieved in this assay reflects that of constitutively dimerized BRAF proteins in vivo, we compared the activities of Tni-FNL–produced autoinhibited monomeric BRAF:14-3-3ζ:MEK complexes with that of dimeric BRAF_2_:14-3-3ζ complexes (SI Appendix, Fig. S9*A*). As shown in Fig. 5*A*, the activity of the autoinhibited BRAF complexes incubated with KRAS and 30% PS liposomes was comparable to that of the dimerized BRAF complexes alone and addition of KRAS and PS liposomes had minimal effect on the dimerized complexes. In addition, quantitative mass spectrometry revealed no significant differences in the phosphorylation state of autoinhibited monomeric versus activated dimeric BRAF, and phosphorylation at the activation segment residues T599 and S602 was not detected (SI Appendix, Fig. S9*C* and Dataset S2).

**Fig. 5.**
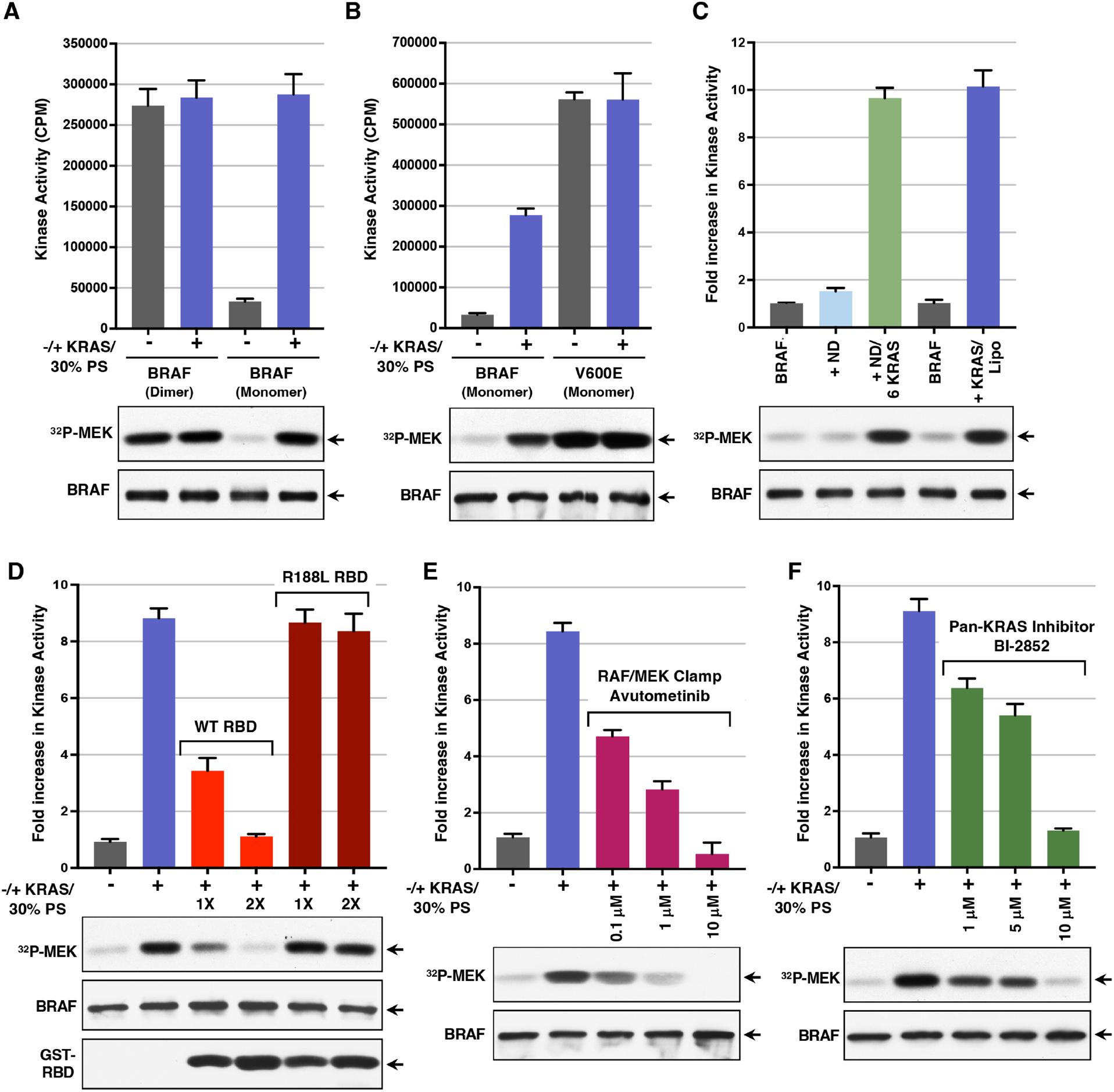
Application of the in vitro BRAF activation assay. *(A)* Tni-FNL-produced BRAF:14-3-3_2_:MEK or BRAF_2_:14-3-3_2_ complexes and were incubated alone or in the presence of GppNHp-KRAS-FMe and 30% PS liposomes for 1 hr prior to monitoring for kinase activity. The graph indicates the average CPM incorporated in kinase-dead MEK from 3 independent experiments ± SD. *(B)* Tni-FNL-produced WT- or V600E-BRAF monomer complexes were evaluated as in *(A*). *(C)* Kinase activity of Tni-FNL BRAF:14-3-3_2_:MEK complexes incubated alone or with lipid nanodiscs is shown. These nanodiscs were either devoid of KRAS or contained 6 KRAS proteins. As a control, BRAF complexes were also incubated with GppNHp-KRAS-FMe and 30% PS liposomes. (*D*) Tni-FNL BRAF:14-3-3_2_:MEK complexes were incubated with GppNHp-KRAS-FMe and 30% PS liposomes in the presence or absence of GST-WT RBD or R188L RBD (at 1 and 2 molar excess) for 1 hr prior to monitoring kinase activity. (*E-F*) Kinase activity of Tni-FNL BRAF:14-3-3_2_:MEK complexes were incubated with GppNHp-KRAS-FMe and 30% PS liposomes, in the presence or absence of the RAF/MEK inhibitor avultometinib (*E*) or the KRAS inhibitor BI-2852 (*F*), as indicated. (*C-F*) The graphs represent the average fold increase in kinase activity, with BRAF activity alone set at 1, based on data from 3 independent experiments ± SD. (*A-F*). Also shown are autoradiographs of the ^32^P-labeled MEK and immunoblot analyses of BRAF levels from representative experiments. For these assays, KRAS-FMe was at a 7 molar excess and 30% PS liposomes at a concentration of 83 μM.

These data support a model in which activation of wild-type BRAF is driven by relief of autoinhibition and dimerization. In contrast, the oncogenic V600E BRAF mutant functions as an activated monomer, with the V600E mutation disrupting autoinhibition and stabilizing the active conformation of the kinase domain (13, 36). Consistent with these reports, we found that the activity of V600E-BRAF was approximately two-fold higher than that of WT BRAF activated in vitro, and nearly 20-fold higher than the basal activity of autoinhibited BRAF complexes (Fig. 5*B*).

Next, to more accurately simulate conditions at the plasma membrane, GppNHp-loaded KRAS was tethered to lipid nanodiscs containing 30% PS, and the RAS-containing nanodiscs were then utilized in the in vitro BRAF activation assay (SI appendix, Fig. S10*A*). We found that incubation of the Tni-FNL BRAF:14-3-3_2_:MEK complexes with lipid nanodiscs alone caused minimal increase in BRAF activity. In contrast, incubation with lipid nanodiscs containing six KRAS proteins had a strong activating effect and was comparable to that observed when KRAS and 30% PS liposomes were added (Fig. 5*C*). These findings demonstrate the ability of KRAS to mediate the in vitro activation of BRAF both in solution and when attached to a membrane-like, lipid matrix.

Finally, we investigated whether the in vitro BRAF activation assay could be used to detect compounds or inhibitors that prevent RAS-mediated BRAF activation. For these studies, we first tested isolated RBD proteins—either wild-type (WT) or those containing the arginine 188 to leucine (R>L) mutation, which disrupts RAS:RBD binding. These proteins were added to the incubation reaction with the autoinhibited BRAF complexes, KRAS, and 30% PS liposomes. As shown in Fig. 5*D*, WT RBD inhibited BRAF activation in a dose-dependent manner, whereas the R188L mutant had no effect. These findings are consistent with the model that the WT RBD, but not the R188L mutant, can compete for KRAS binding, thereby reducing KRAS’s ability to interact with the autoinhibited complexes and promote BRAF activation.

We next examined the effect of avutometinib, a compound that stabilizes BRAF in the autoinhibited conformation by functioning as a RAF/MEK clamp. As shown in Fig. 5*E*, stabilization of the autoinhibited BRAF complex with avutometinib treatment could effectively prevent BRAF activation in vitro.

To further confirm that BRAF activation in this assay is RAS-dependent, we examined the effect of BI-2852, a KRAS inhibitor that binds to the switch I/II pocket of both GDP and GTP-bound KRAS, thereby blocking interactions with SOS and downstream effectors such as RAF (37). Consistent with this mechanism of action, BI-2852 inhibited BRAF activation in a dose-dependent manner, with the 10 µM dose effectively blocking BRAF activation (Fig. 5D). Similar results were also observed with the KRAS inhibitor, MRTX1133 (SI appendix, Fig. S10*B*) (38), further confirming the importance of KRAS in relieving BRAF autoinhibition in this system.

## Discussion

To further elucidate the BRAF activation process, we developed an in vitro BRAF activation assay utilizing autoinhibited BRAF:14-3-3_2_:MEK complexes and other purified components. Our results are consistent with the model that binding interactions between the BRAF RBD and KRAS can initiate the disassembly of the autoinhibited complex (Fig. 6). In vitro, KRAS-FMe alone promoted BRAF activation in a dose-dependent manner, which was further augmented by the presence of PS-containing liposomes. The activating effect of KRAS required that KRAS be in its active state and was dependent on BRAF’s ability to dimerize. Although lipid nanodiscs alone had minimal effect on BRAF activity, lipid nanodiscs containing tethered active-state KRAS proteins also mediated BRAF activation, with the levels of activity being comparable to those obtained with soluble KRAS and 30% PS liposomes. Moreover, the level of activity achieved in the in vitro BRAF activation assay was similar to that of BRAF proteins that constitutively dimerize in vivo. Further confirming the importance of RBD:KRAS binding, inhibitors that disrupt this interaction could suppress KRAS-mediated BRAF activation in a dose-dependent manner. Specifically, isolated RBD proteins competent to bind RAS and a KRAS inhibitor targeting the Switch I/II pocket to block effector interactions, both effectively inhibited BRAF activation in vitro. Thus, these findings indicate that KRAS plays a more active role in promoting BRAF activation than merely serving as a membrane tether. However, as the autoinhibited complex disassembles, the presence of the plasma membrane allows the CRD to interact with negatively charged lipids, further promoting the exposure of the kinase domain for dimerization and activation.

**Fig. 6.**
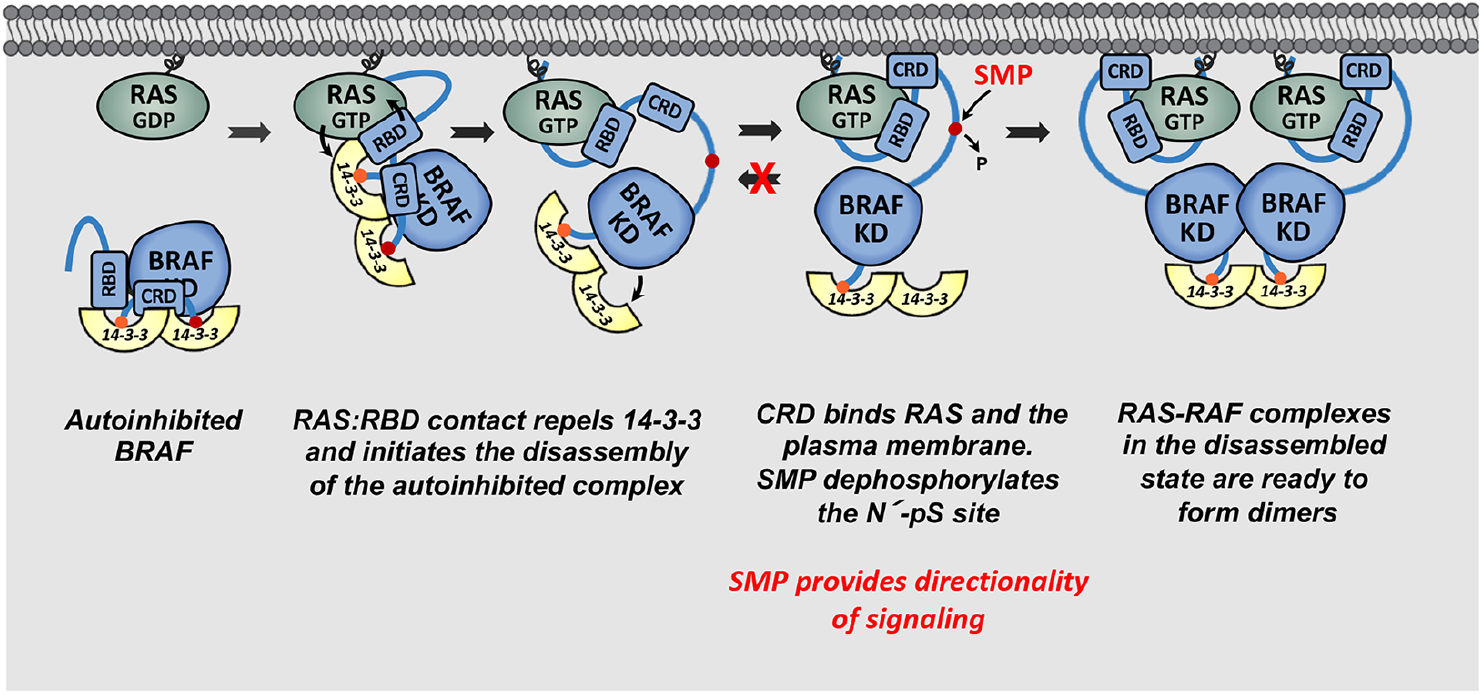
Model for RAS-driven BRAF activation. See text for details.

Notably, the SMP phosphatase complex, which forms at the plasma membrane in response to signaling events and dephosphorylates the N’ pS365 14-3-3 binding site, had only a small effect on BRAF activity levels under the in vitro assay conditions. However, SMP was required for pS365 dephosphorylation, and this process was most efficient when both KRAS and PS-containing liposomes were present, indicating that these conditions increase the exposure or accessibility of the pS365 site. Although SMP had minimal effect on BRAF catalytic activity in vitro, its function may be critical in live cells, where multiple effector cascades are simultaneously activated in response to signaling events. In this context, SMP-mediated dephosphorylation of pS365 may play an important role in promoting the forward directionality of signaling by preventing the rebinding of 14-3-3 and facilitating dimerization of disassembled BRAF monomers (Fig. 6). Consistent with this model, we observed a more pronounced effect of SMP on BRAF activity when KRAS was at suboptimal levels. These findings suggest that under conditions of constitutive oncogenic KRAS signaling, where both positive and negative feedback loops are engaged, or under conditions of low KRAS activity, sustained dephosphorylation of pS365 may be necessary to ensure that 14-3-3 remains unbound, thereby enabling RAF dimer formation and signal propagation.

In addition, our study provides insights regarding the differences observed for autoinhibited BRAF:14-3-3_2_:MEK complexes isolated from Sf9 insect cells and human 293FT cells. As previously reported, we found that the Sf9-derived complexes exhibited a high basal activity that increased over time during the incubation period, suggesting a less stable autoinhibited state. In contrast, we found that BRAF:14-3-3_2_:MEK complexes produced in Tni-FNL insect cells were similar in structure and activity to those produced in 293FT cells. Specifically, the RBD was resolved in the Tni-FNL structure, but not in the Sf9 structure, and formed a similar interface with 14-3-3. Moreover, like the 293FT complexes, the Tni-FNL complexes had a low basal activity that remained unchanged throughout the incubation reaction but were highly activatable when incubated with KRAS and 30% PS liposomes or with KRAS-tethered lipid nanodiscs. Although we did not assess in detail the factors contributing to the differences between the Sf9 and Tni-FNL complexes, it is possible that the signaling state of the cells may play a role, as the Sf9 complexes displayed higher levels of phosphorylation, particularly at known ERK-dependent feedback sites, which could affect their stability. Further mass spectrometry analyses of Tni-FNL produced autoinhibited BRAF monomers and activated BRAF dimers revealed no significant differences in the phosphorylation state of the Tni-FNL produced complexes, and that phosphorylation on the activation segment residues T599 and S602 was not detected. These findings support the model that activation of wild type BRAF is mediated by relief of autoinhibition and RAF dimerization rather than phosphorylation of the activation segment.

Finally, our study demonstrates that the in vitro BRAF activation assay is effective in detecting proteins and compounds that inhibit the BRAF activation process. Thus, this assay represents a reliable platform for future research and drug screening initiatives aimed at targeting BRAF. Specifically, this assay can identify agents that disrupt BRAF catalytic activity or prevent critical interactions, as well as those that stabilize the autoinhibited state, representing a novel strategy for inhibiting BRAF function.

## Methods

### In Vitro BRAF Activation Assay

For the BRAF activation assay, we used purified autoinhibited BRAF complexes (BRAF:14-3-3ζ:MEK) that had been assessed by quantitative mass spectrometry to confirm high stoichiometry of phosphorylation at the regulatory pS365 and pS729 sites. The autoinhibited BRAF complexes (250 nM) were incubated in the presence or absence of purified full-length KRAS4B-FMe proteins, PS-containing liposomes, SMP complexes, and/or combinations thereof. Reactions were carried out for 1 hour at 24 °C in a buffer containing 30 mM Tris, pH 7.4, in a final volume of 12 μL. In most assays, KRAS was added at a 7-fold molar excess relative to the BRAF complexes, SMP at a 1.2-fold molar excess, and liposomes at a final concentration of 83 μM. In certain experiments, KRAS-tethered nanodiscs were used in place of purified KRAS proteins and liposomes. When small-molecule inhibitors were tested, they were added at the start of the incubation reaction.

To determine the kinase activity of BRAF after the activation reaction, 38 μl of kinase buffer was added to each sample to give a final concentration of 30 mM Tris, pH7.4, 10 mM MnCl_2_, 1 mM dithiothreitol, 150 μM ATP, 5 mM MgCl_2_. 20 μCi [γ-^32^P]ATP and 0.1 mg of purified kinase-inactive MEK, and samples were incubated for 30 minutes at 24°C. Reactions were terminated by the addition of boiling 4x sample buffer (33% glycerol, 0.3 M DTT, 6.7% SDS, 0.1% bromophenol blue), and proteins were resolved by SDS-PAGE and transferred to 0.2 μm nitrocellulose membranes. Protein levels were confirmed by western blot analysis. The amount of radioactivity incorporated into the kinase-inactive MEK was visualized by autoradiography and quantified by scintillation counting.

Additional experimental procedures are in *SI Appendix* Materials and Methods. Reagents and resources are listed in *SI Appendix* Table S1.

## DATA Availability

The cryo-EM density maps were deposited in the Electron Microscopy Data Bank (EMDB) with the accession code EMD-70053 for the BRAF:14-3-3_2_:MEK global refinement map. The corresponding atomic model for the complex was deposited in the Protein Data Bank (PDB) under the accession code 902Z. All simulation parameter files and select simulation files will be made available at https://bbs.llnl.gov/. *These structures will be deposited and an accession code obtained upon acceptance.

## Acknowledgements

We thank Vanessa Wall, Nicholas Wright, Hannah Ambrose, Ashley Mitchell, Kayla Russell, and Katie Reeley for protein purification. This project was funded in part with federal funds from the National Cancer Institute (NCI), National Institutes of Health (NIH) under Project numbers ZIA BC 010329 to DKM and ZIA BC 011744 to PZ and under Contract 75N91019D00024 to Frederick National Laboratory for Cancer Research (FNLCR). This work utilized the cryoEM-Facility in the Center for Structural Biology, NCI at Frederick, as well as computational resources of the Frederick Research Computing Environment cluster. Part of this work was also performed under the auspices of the US Department of Energy (DOE) by Lawrence Livermore National Laboratory under Contract DE-AC52-07NA27344. This work has been supported by the NCI-DOE Collaboration established by the US DOE and the NCI/NIH. This research utilized resources of the Oak Ridge Leadership Computing Facility (OLCF), which is a DOE Office of Science User Facility supported under Contract DE-AC05-00OR22725. For computing time, the authors thank the Livermore Institutional Grand Challenge for time on Lassen and the Advanced Scientific Computing Research Leadership Computing Challenge (ALCC) for time on Summit. For computing support, the authors thank LC and OLCF staff. The contributions of the NIH author(s) were made as part of their official duties as NIH federal employees, are in compliance with agency policy requirements, and are considered Works of the United States Government. However, the findings and conclusions presented in this paper are those of the author(s) and do not necessarily reflect the views of the NIH or the U.S. Department of Health and Human Services. Release: LLNL-JRNL-XXXXXX.

