## Supplemental information for "An In Vitro BRAF Activation Assay Elucidates Molecular Mechanisms Driving Disassembly of the Autoinhibited BRAF State"

**This PDF file includes:**

- Supplementary Materials and Methods
- Figs. S1 to S10 and Tables S1 to S2
- Captions for Movies S1 to S2 and Datatset S1 and S2.
- References for SI reference citations

**Other supplementary materials for this manuscript include the following:**

- Movies S1 to S2
- Dataset S1 and 2

**Supplementary Materials and Methods**

**A list of Reagents and resources can be found in Table S1.**

**Molecular Dynamics Simulations.** All-atom molecular dynamics (MD) simulations were performed on the BRAF:14-3-3 complex, utilizing the CHARMM36m force field and based on the cryo-EM structures PDB ID: 7MFD and PDB ID: 6NYB. The recently developed protein model for 7MFD was used in these simulations (1). In brief, the BRAF:14-3-3$_2$ complex was constructed from PDB ID: 7MFD with several modifications. ATP, $Mg^{2+}$, and zinc ions were added, MEK was removed, and missing residues for 14-3-3 and BRAF (encompassing residues 156 to 738) were modeled using MOE. The protein ends were capped, and BRAF S365, S446, and S729 were added in their phosphorylated states. The larger intrinsically disordered regions—Loop I (residues 274 to 359) and Loop II (residues 371 to 448)—were generated in multiple configurations, resulting in an ensemble of different loop conformations for the starting structures. A total of 27 different loop configurations were selected, and five independent replicas were run for a minimum of 500 ns each, resulting in 135 simulations aggregating over 73 μs of simulation data.

The 6NYB model was built following the same strategy as the 7MFD model, with the addition of modeling in the missing RBD. The RBD from 7MFD was used for its internal configuration, and seven different RBD starting conformations were used. The RBD was positioned in the following ways: 1) aligned to RBD-CRD of 7MFD, 2) aligned to RBD-CRD of the RBD-CRD X-ray structure PDB ID:6XI7 (2), where the RBD is rotated by ~180° compared to 7MFD, so intermediate rotations were also explored, 3) the configuration of #2 rotated by 60°, 4) the configuration of #2 rotated by 120°, 5-7) three additional RBD configurations were selected based on a previous RBD-CRD simulation campaign (3). Several configurations of Loop I and Loop II were constructed for each RBD placement with five independent replicas run for each, resulting in a total of 638 simulations. Most simulations were run for 500 ns, aggregating over 406 μs of simulation data.

The first models before final Loop and RBD placements were set up using CHARMM-GUI (4). The following topology construction and minimization were performed with GROMACS (v2019.6) (5), followed by format conversion using the ParmEd gromber tool in

AmberTools 19. These CHARMM36m simulations were run with one protein in a water box of dimensions 14.5 nm x 14.5 nm x 14.5 nm with 0.15 M NaCl, using the TIP3P water model (6), with water constraints applied via SETTLE (7). All simulations were run with a time step of 2 fs for a minimum of 500 ns using AMBER18 with mixed-precision (SPFP (8)).

Hydrogen bonds were identified by employing the HBonds plugin within VMD (9). A distance threshold of 3.5 Å and an angular cutoff of 30° were set as criteria for defining hydrogen bonds. Specifically, hydrogen bonds were considered present if the donor-acceptor distance was less than or equal to 3.5 Å and the hydrogen-donor-acceptor angle was less than or equal to 30°. All simulation visualization and density plots were created with VMD.

**Generation of Baculovirus Constructs.** Gateway Entry clones for human BRAF amino acids 2-766 (WT or V600E) and 438-766, as well as human MAP2K1 (MEK1) amino acids 1-393, were synthesized by ATUM, Inc. and optimized for expression in insect cells. All constructs contained an upstream tobacco etch virus (TEV) protease cleavage site (ENLYFQ/G). Gateway LR recombination was then used to transfer the BRAF Entry clones to pDest-636 (Addgene #159574) to generate a baculovirus expression clone with a His6-MBP (maltose binding protein) N-terminal fusion tag. Similarly, the MAP2K1 Entry clone was transferred to pDest-635 (Addgene #159582) to generate a baculovirus expression clone with a His6 N-terminal fusion tag. BRAF expression clones were used to generate bacmid DNA in *E. coli* DE105 (10), which contains a baculovirus genome that co-expresses human CDC37 and YWHAZ (14-3-3ζ) proteins. In contrast, MAP2K1 expression clones were used to generate bacmid DNA in E. coli DE95 (11).

**Preparation of BRAF Complexes.** Mammalian BRAF:14-3-$3_2$:MEK complexes were prepared as previously described (12). Briefly, forty confluent 10 cm dishes of 293FT cells stably expressing Halo-tagged $BRAF^{WT}$ were washed twice with 5 mL cold PBS and lysed in Triton lysis buffer (1% Triton X-100, 137 mM NaCl, 20 mM Tris pH 8.0, 0.15 U/mL aprotinin, 1 mM phenylmethylsulfonyl fluoride, 20 μM leupeptin; 0.5 mM sodium vanadate; 1mL per 10 cm dish) at 4°C for 15 minutes on a rocking platform. Lysates were collected and clarified of debris by centrifugation at 14,000 rpm for 10 minutes at 4°C. Five mL of Halolink resin (Promega) washed twice in Triton X-100 lysis buffer was then added to the clarified supernatant and incubated at

4°C for 2 hours on a rocking platform. Beads containing the bound BRAF complexes were washed twice with Triton X-100 lysis buffer and three times with elution buffer (137 mM NaCl and 20 mM Tris pH 8.0). The bead-bound complexes were then resuspended in 2.5 mL elution buffer containing 50 μL Halo-TEV (Promega) and incubated at 4°C for 2 hours on a rocking platform. The beads were then pelleted, and the supernatant containing the eluted BRAF complexes was applied to a Superose 6 Increase 10/300 GL column (Cytiva) pre-equilibrated with a buffer containing 20 mM Tris pH 8.0 and 137 mM NaCl. Proteins from the peak fraction corresponding to BRAF:14-3-$3_2$:MEK complexes were collected and analyzed by SDS-PAGE and silver staining.

For the production of BRAF complexes in insect cells, recombinant baculoviruses were generated and titered as previously described (13). Briefly, Sf9-produced BRAF:14-3-$3_2$:MEK complexes were prepared by infecting Sf9 cells with a recombinant baculovirus encoding His6-tev-Hs.MAP2K1 as well as a baculovirus encoding His6-MBP-tev-Hs.BRAF(2-766), Hs.CDC37, and Hs.14-3-3ζ with multiplicity of infection (MOI) of 1 and 2, respectively. The infected cells were then incubated at 27°C with shaking for 72-80 hours until the viability dropped below 70% as determined by Trypan Blue exclusion dye. Cells were then pelleted by centrifugation at 1200 x g for 15 minutes.

Tni-FNL-produced monomeric BRAF:14-3-$3_2$:MEK complexes were prepared by infecting Tni-FNL cells with the same recombinant viruses as above with equal MOI of 3. For Tni-FNL-produced dimeric $BRAF_2$:14-3-$3_2$ complexes and monomeric V600E-BRAF:14-3-$3_2$ complexes, cells were infected with the baculovirus encoding His6-MBP-tev-Hs.BRAF (2-766) WT or V600E, Hs.CDC37 and Hs.14-3-3ζ. The Tni-FNL cells were then incubated at 21°C with shaking for 72-80 hours until the viability dropped below 70%. Cells were then pelleted by centrifugation at 1200 x g for 15 minutes.

The pellets were then lysed in a Microfluidics LV1 in 50 mM Tris-Cl pH 8.5, 500 mM NaCl, 1 mM TCEP, 5 mM $MgCl_2$, protease inhibitor cocktail (1:200 dilution, P8849, Sigma Aldrich) and clarified via ultracentrifugation (100,000 × *g)* for 30 minutes. The clarified lysates were applied to a 5 ml HiTrap Nickel HP column equilibrated previously in lysis buffer and eluted with a 20-column volume (CV) gradient from 0-100% equilibration buffer with 500 mM imidazole, collecting 5 ml fractions. Fractions were assayed via SDS-PAGE, and fractions containing BRAF complexes were pooled and digested with His6-TEV, while dialyzing against

50 mM Tris-HCl pH 8.5, 500 mM NaCl, 1 mM TCEP, 5 mM $MgCl_2$. Following dialysis, the sample was applied to a 5 ml HiTrap Nickel HP column to remove the remaining affinity tags and TEV. The flowthrough was collected, concentrated via tangential flow filtration and applied to a Superdex 200 16/60. Fractions collected from the Superdex 200 were analyzed by SDS-PAGE, and fractions containing the BRAF complexes were pooled and concentrated to 1 mg/ml via Amicon centrifugal concentrators. Aliquots of the purified complexes were snap frozen in liquid nitrogen and stored at -80$^{o}$C.

**Production of KRAS proteins and the SMP phosphatase complex.** Wild-type and D154Q-KRAS4b (2-185)-FMe and C118S-KRAS4b (1-185) were produced and purified following previously published protocols (14). SMP complexes were generated and purified as described in Snead et al. (13).

**GppNHp loading of KRAS proteins.** KRAS-FMe and KRAS C118S (1-185) were diluted to a final concentration of 50 μM in 40 mM Tris pH 7.4, and 0.1 mM $ZnCl_2$. Ammonium sulfate was added to a final concentration of 500 mM, followed by the addition of 2.5 mM GppNHp and 1 mM Alkaline Phosphatase beads. The solution was allowed to rotate at room temperature for 1 hour, centrifuged to pellet the beads and the supernatant was removed. Magnesium chloride was added to the supernatant to a final concentration of 20 mM and the protein solution was rotated at room temperature for 1 hour. Excess nucleotide was removed using a HiPrep 26/10 Desalting column equilibrated with 20 mM HEPES pH 7.0, 150 mM NaCl, and 1.5 mM $MgCl_2$.

**Preparation of Liposomes.** Liposomes composed of 70 POPC:30 POPS were produced at a lipid concentration of 1 mM total lipid in 1 mL of 20 mM HEPES pH 7.4, 150 mM NaCl or 0.76 mg/mL lipid. Lipids dissolved in chloroform were transferred to a glass vial and dried under inert gas before being placed in a lyophilizer for 2 hours to ensure complete dryness. After removal from the vacuum chamber the lipids were allowed to hydrate for 1 hour in HEPES buffered saline. The lipids were briefly vortexed and sonicated to ensure the lipid film lifts from the glass. The turbid solution was transferred to a mini-extruder (Avanti Polar Lipids, Cat# 610000-1EA) set with a 100 nm pore polycarbonate membrane and extruded 10 times. Liposomes containing an eight-lipid mixture of cholesterol, phosphatidylcholines (1-palmitoyl-2-oleoyl-*sn*-glycero-3-

phosphocholine, POPC and 1-palmitoyl-2-arachidonoyl-*sn*-glycero-3-phosphocholine, PAPC), phosphatidylethanolamines (1-palmitoyl-2-oleoyl-*sn*-glycero-3-phosphoethanolamine, POPE and 1,2-dilinoleoyl-*sn*-glycero-3-phosphoethanolamine, DIPE), phosphatidylserine (1-palmitoyl-2-arachidonoyl-*sn*-glycero-3-phospho-L-serine, PAPS), phosphatidylinositol bisphosphate (1-stearoyl-2-arachidonoyl-sn-glycero-3-phospho-1'-myo-inositol-4',5'-bisphosphate, PIP2), and sphingomyelin (N-palmitoyl-D-erythro-sphingo-sylphosphorylcholine, DPSM) were prepared as previously described (15, 16).

**Preparation of Lipid Nanodiscs and RAS-containing Lipid Nanodiscs.** Lipid nanodiscs were produced to effectively tether two different stoichiometric amounts of KRAS, 65.6 POPC:30 POPS:4.4 PEMCC (4.4 PEMCC) and 60 POPC:30 POPS:10 PEMCC (10 PEMCC). 29.47 mg and 28.66 mg of total lipid were dried in the same manner as described above for the 4.4 PEMCC and 10 PEMCC lipid mixtures respectively. After drying, the lipids were resuspended in 600 µL of 20 mM HEPES (pH 7.0), 150 mM NaCl, 130 mM cholate buffer for a final concentration of 65 mM total lipid. Samples were vortexed and sonicated as needed to ensure all lipids were suspended. The lipids were mixed with the nanodisc belt protein MSP1E3D1 at a ratio of 135:1 lipid to protein and rotated at room temperature for 1 hour. Bio-Beads SM-2 Absorbent Media purchased from Bio-Rad Laboratories (Hercules, CA, USA) were added to the lipid-protein mixture to remove the detergent and allowed to rotate overnight at 4°C.

The next day the Bio-Beads were removed via centrifugation through a large pore filter and the nanodiscs were passed over a Superdex 200 Increase 10/300 GL size exclusion column (Cytiva) equilibrated with a buffer composed of 20 mM HEPES (pH 7.0) and 150 mM NaCl. Nanodisc size was determined by checking fractions of the major peak via Dynamic Light Scattering (DLS). DLS measurements were performed on a DynaPro PlateReader-II from Wyatt Technology (Santa Barbara, CA, USA). Fractions were collected with an average size of 13.2 ± 0.2 nm and concentrated for tethering with RAS. The nanodisc preparations were mixed with GppNHp-loaded KRAS C118S (1-185) at a molar ratio of 3:1 RAS to lipid nanodisc and rotated overnight. The next morning the KRAS-nanodiscs were quenched with beta-mercaptoethanol for 1 hour. The solutions of RAS-nanodiscs were passed through the same size exclusion column equilibrated with 20 mM HEPES (pH 7.0), 150 mM NaCl, and 1.5 mM $MgCl_2$.

Fractions of RAS-nanodisc were checked via the ReFeyn Two MP mass photometer (Waltham, MA, USA) to determine the stoichiometry of RAS to nanodisc. Fractions containing stoichiometric amounts of 6 KRAS per nanodisc were collected and concentrated. The concentration of RAS was calculated from the absorbance for each RAS-nanodisc. This determination was made by multiplying the absorbance reading with the fraction of each molar extinction coefficient, ε, contribution and then multiplying the ε for the stoichiometric amount of the protein (e.g. 6 KRAS molecules per 2 molecules of MSP1E3D1 nanodisc belt protein).

The concentration of lipid per nanodisc was determined using the contribution of absorbance from the MSP1E3D1 per sample and the number of lipids composing the nanodisc. The number of lipids per nanodisc were determined by calculating the surface area of both faces of the nanodisc and dividing by the surface area of the phospholipid head group.

Contribution of RAS in RAS-nanodisc:

$$\frac{\varepsilon_{RAS}\ x\ \#RAS}{(\varepsilon_{RAS}\ x\ \#RAS) + (\varepsilon_{MSP1E3D1}\ x\ 2)}$$

Where the #RAS is expected to be either 3 or 6 depending on the sample chosen. This fraction is then multiplied by the overall concentration of the RAS-nanodisc to determine the concentration of RAS.

Contribution of nanodisc belt protein MSP1E3D1 and concentration of lipid:
To determine the concentration of total lipid the first step is to determine the concentration of nanodisc.

$$\frac{\varepsilon_{MSP1E3D1}\ x\ 2}{(\varepsilon_{RAS}\ x\ \#RAS) + (\varepsilon_{MSP1E3D1}\ x\ 2)}$$

The fraction is multiplied by the concentration of RAS-nanodisc to give the concentration of nanodisc, which is then multiplied by the number of lipids per nanodisc.

$$\#\ lipid\ per\ nanodisc = 2\left(\frac{\pi r^2}{0.64}\right)$$

Where a single face of the nanodisc is calculated using the area of a circle divided by the surface area of a phospholipid head group (0.64 $nm^2$).

**Peptide Reaction Monitoring Mass Spectrometry Analysis.** BRAF:14-3-$3_2$:MEK complexes were precipitated with ice-cold acetone at -20°C overnight. Acetone was decanted then samples were resuspended in 8 M urea in 100 mM ammonium bicarbonate for proteolytic digestion (17, 18). DTT (dithiothreitol) was added to reduce disulfide bonds for 1 hour. IAA (iodoacetamide) was added to alkylate free cysteines for 45 minutes in the dark then quenched with DTT. Samples were further diluted in 100 mM ammonium bicarbonate to lower urea concentration. Proteins were digested overnight at 37°C with 500 ng of AspN, GluC, or Trypsin digestion enzymes. Digestion was quenched with concentrated formic acid. Peptides were desalted using Pierce C18 Spin columns (Thermo Fisher Scientific). Eluted peptides were evaporated to dryness using a SpeedVac, then resuspended in 0.2% formic acid and transferred to autosampler vials. AspN, GluC, or Trypsin digested protein samples were analyzed by liquid chromatography and subsequent tandem mass spectrometry (LC-MS/MS on an UltiMate 3000 chromatographic system (Thermo Fisher Scientific) with reverse-phase separation, using an Acclaim™ PepMap™ 100 C18 HPLC trap column (5 µm, 100 Å, 0.1 x 20 mm, Thermo Fisher Scientific) and Acclaim™ PepMap™ 100 C18 HPLC analytical column (3 µm, 100 Å, 0.075 x 500 mm, Thermo Fisher Scientific), coupled to an Orbitrap Fusion Lumos mass spectrometer. Each digested sample was injected in triplicate for LC-MS/MS analysis by either higher-energy collisional dissociation (HCD), electron transfer dissociation with supplemental HCD (EThcD), or collision-induced dissociation (CID). All files were searched in Proteome Discoverer version 3.1.1.93 (Thermo Fisher Scientific) and analyzed manually in Freestyle 1.5.93.34 (Thermo Fisher Scientific), XCalibur Qual Browser version 4.2.47 (Thermo Fisher Scientific), and Skyline version 24.1.0.414 (University of Washington). Graphical fragment maps were generated using the Xtract algorithm in Xcalibur and ProSight Lite (19).

**Cryo-EM Data Acquisition, and Processing.** After gel filtration, the Tni-FNL-produced BRAF:14:3-$3_2$:MEK complexes (in 20 mM Tris pH 8.0 and 137 mM NaCl) were concentrated to 0.18 - 0.20 mg/mL by centrifugation using Pall's Microsep™ advance centrifugal device with a 30K pore size. The samples were supplemented with NaCl and DTT to reach final concentrations of 200 mM and 10 mM, respectively, before freezing. Quantifoil Au R1.2/1.3 holey carbon grids, 300 mesh, were subjected to glow discharge for 45 seconds at 25 mA in an alcohol environment. A volume of 1.5 µL of protein solution was applied to each side of the grids, and the samples were vitrified using a Leica EM GP2 plunge freezer with a blotting time of 2.5 seconds at 4 °C and humidity above 85%. The grids were subsequently plunge-frozen into liquid ethane cooled to approximately -180 °C by liquid nitrogen.

Cryo-EM data acquisition was conducted at the cryo-EM facility in the Center for Structural Biology, NCI-Frederick, using a Talos Arctica G2 (Thermo Fisher) equipped with a Gatan K3 direct detector and energy filter, operated at 200 keV. The slit width of the energy filter was set to 20 eV. Data were collected in super-resolution mode at a nominal magnification of 100,000×, corresponding to 0.405 Å/pixel. 50 frames per movie were acquired for a total dose of approximately 50 elections/Å2. Then, using EPU data collection software (Thermo Fisher), the micrographs were dose-fractionated into 50 frames for a total electron exposure of 50 electrons per $Å^2$, with defocus values ranging from -0.8 to -2.5 μm. A total of 39,636 micrographs were collected.

All cryo-EM data analysis was done using Cryosparc 3.3 (20). Movies were imported, patch-motion and patch-CTF corrected. Movies were binned to the physical pixel size in the patch motion step. The micrographs were screened to remove low-quality images and those with unqualified CTF power spectra. An initial subset of particles was picked using blob picker and used to train a Topaz(21, 22) model that was used to pick particles in the entire data set. Particles were curated using multiple rounds of 2D classifications, after which duplicated particles were removed. An initial 3D classification with Ab-initio and heterogeneous refinement (23) identified a volume corresponding to BRAF:14-3-$3_2$:MEK. Further non-uniform refinement achieved a map with 3.8 Å resolution, as determined by the gold standard FSC. A graphical summary of the cryo-EM data processing is presented in Supplementary Fig. S7*E*. Cryo-EM reconstruction statistics are detailed in Supplementary Table S2. The refined maps were deposited in the EMDB database. The BRAF:14-3-$3_2$:MEK complex model was built by the rigid-body fitting of the entire complex (PDB ID: 7MFD) into the global refined map. Masked-based local refinement was also performed on different parts of the structure, which resulted in no further improvement of the map. Fitting of the models into the map was initially done using UCSF Chimera (24). Manual adjustment of the model was performed in Coot (25), followed by iterative rounds of real space refinement in Phenix (26) and manual fitting in Coot. Model validation was done using statistics from Ramachandran plots and MolProbity scores in Phenix and Coot (27, 28). Statistics for the final refinements are shown in Supplementary Table S2. Figures were generated by UCSF Chimera. Structure deviations and electrostatic potential of surfaces were calculated using

MatchMaker and Columbic Surface plugins, respectively, in UCSF Chimera. The final model was deposited in the PDB database.

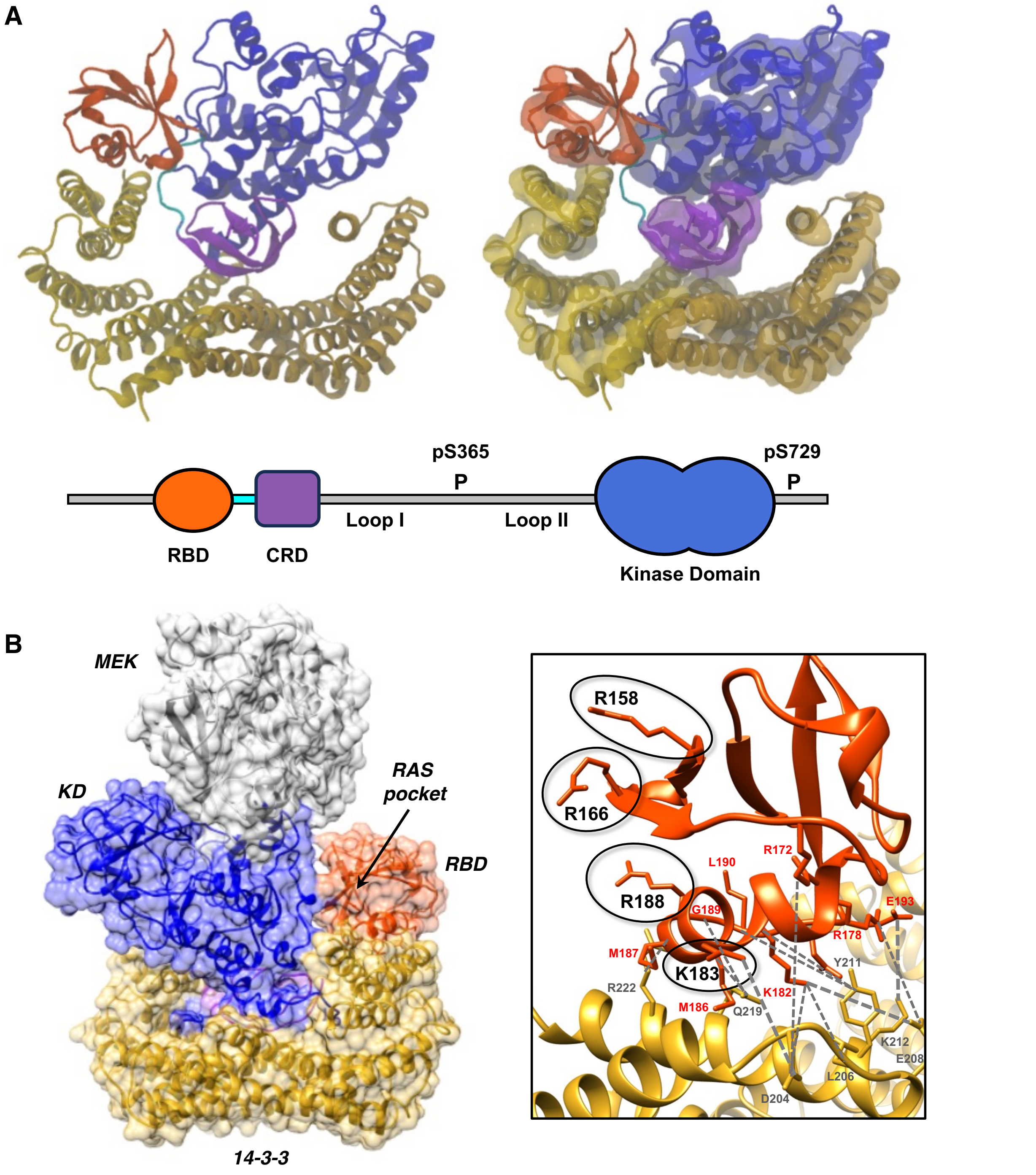

**Fig. S1.** RAS-binding interface of the BRAF RBD. (*A*) To assess the stability of the RBD:14-3-3 interface, 135 independent all-atom simulations were run with 27 different configurations of the unstructured loop I and II regions. A snapshot from one of the simulations after initial equilibrium shows the relative orientation of the different domains (left), with the RBD in red, the RBD-CRD loop in cyan, the CRD in purple, the KD in blue, and the 14-3-3 proteins in yellow/gold. For clarity, solvent molecules, ATP, $Zn^{2+}$ ions, and the unstructured loops are excluded. A volume density map of the protein domains (isovolume = 0.5), averaged over all 135 simulations, is aligned with the reference structure (right). The RBD shows good density for most of the structure, indicating high stability and positional consistency. A schematic depiction of BRAF is also shown for reference. (*B*) Cryo-EM structure of the autoinhibited BRAF:14-3-$3_2$:MEK complex is shown indicating the "RAS pocket" for RAS:RBD binding (left). Basic residues of the BRAF-RBD (R158, R166, K183, and R188) that form critical ionic bonds with the switch I domain of RAS are indicated on the right. Hydrogen bonds observed between RBD and 14-3-3 residues in the molecular dynamics simulations of the autoinhibited BRAF complex are also shown as dashed gray lines.

**A**

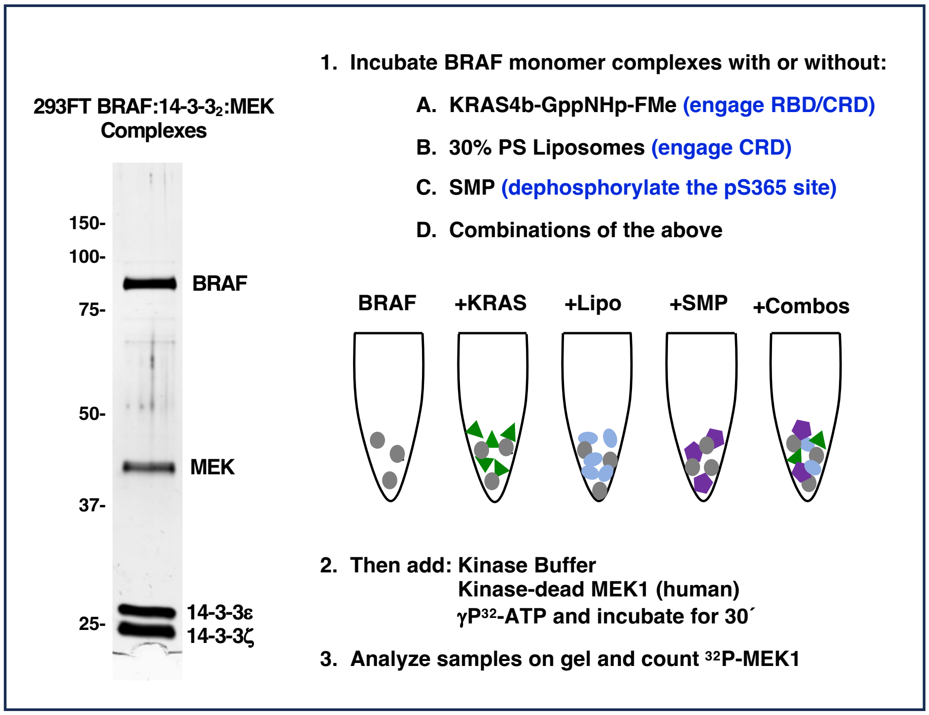

**Assay specifically monitors changes in BRAF activity that occur during the incubation period

**B**

*Quantitative MS Analysis*

| Residue | Avg. % Phos. | Range |
|---|---|---|
| S365 | 97.83% | 96.70-99.38% |
| S729 | 99.57% | 99.24-99.72% |

**C**

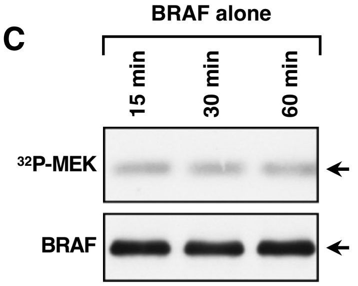

**Fig. S2.** In vitro BRAF activation assay and analysis of 293FT- autoinhibited BRAF:14-3-3$_2$:MEK complexes. (*A*) Workflow for the in vitro BRAF activation assay is depicted (right). Also shown is a silver-stained SDS-PAGE gel of a representative preparation of 293FT-produced autoinhibited BRAF:14-3-3$_2$:MEK complexes used in the in vitro activation assays, with the BRAF, MEK, and 14-3-3 proteins indicated (left). (*B*) Preparations of 293FT-produced autoinhibited BRAF complexes used in the in vitro BRAF activation assays were assessed for phosphorylation at the pS365 and pS729 14-3-3 binding sites by quantitative mass spectrometry. The average stoichiometry of phosphorylation at these sites is shown along with the range of phosphorylation detected across 5 independent preps. (*C*) 293FT-produced autoinhibited BRAF complexes were incubated alone at 25°C for 15, 30, or 60 minutes prior to monitoring for kinase activity in vitro using kinase-dead MEK as a substrate. An autoradiograph of the $^{32}P$-labeled MEK and immunoblot analysis of the BRAF levels are shown, indicating that the basal kinase activity of the autoinhibited BRAF complex remains constant during the incubation reaction.

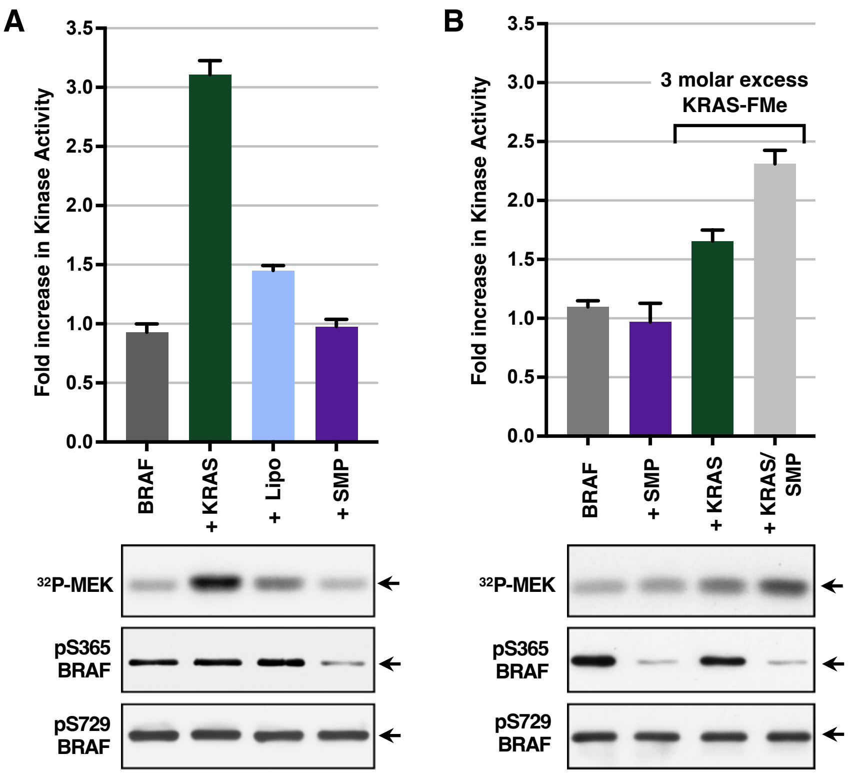

**Fig S3**. Effect of SMP on BRAF activation in vitro (*A*) Autoinhibited BRAF complexes were incubated alone or in the presence of GppNHp-bound KRAS-FMe (7 molar excess), 30% PS-containing liposomes (41μM), or SMP (1.2 molar excess) for 1 hr and then monitored for kinase activity (*E*) Autoinhibited BRAF complexes were incubated alone or with SMP (1.2 molar excess) in the presence or absence of GppNHp-KRAS-FMe (3 molar excess) prior to monitoring for kinase activity. (*D-E*) The graphs represent the average fold increase in kinase activity, with BRAF activity alone set at 1, based on data from 3 independent experiments ± SD. Also shown are autoradiographs of the $^{32}$P-labeled MEK and immunoblot analyses of pS365, pS729, and total BRAF levels from representative experiments.

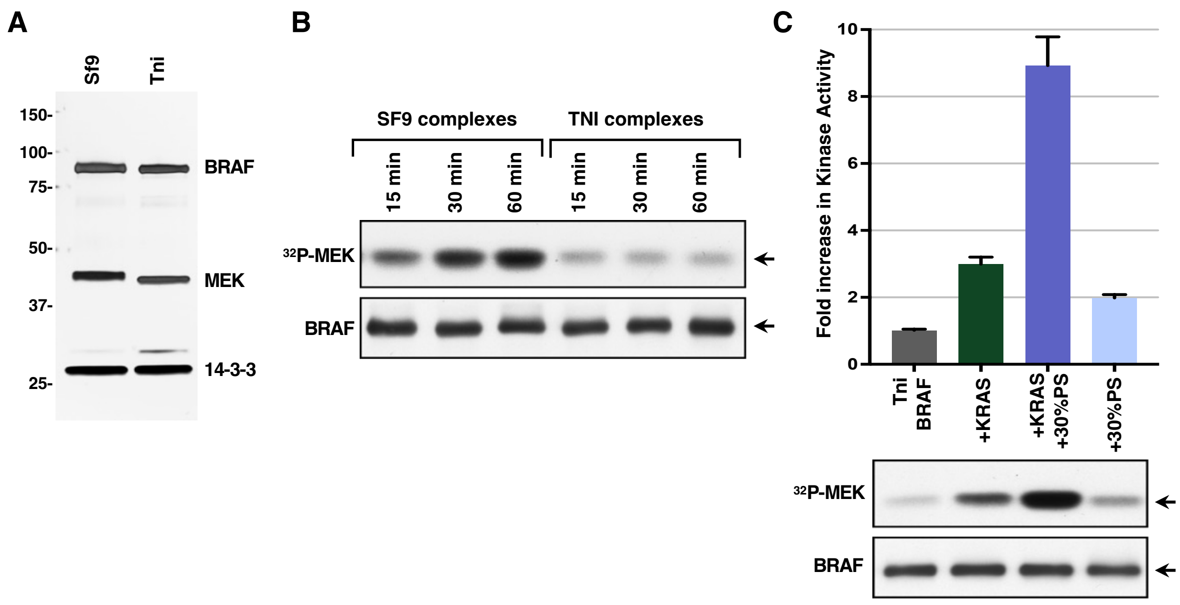

**Fig. S4.** Comparison of Sf9- and Tni-FNL-produced human BRAF:14-3-3$_2$:MEK complexes. (*A*) Complexes were analyzed by SDS-PAGE and silver staining. BRAF, MEK, and 14-3-3ζ are indicated. (*B*) BRAF:14-3-3$_2$:MEK complexes produced in Sf9 and Tni-FNL insect cells were incubated alone at 25°C for 15, 30, or 60 minutes prior to monitoring for kinase activity in vitro using kinase-dead MEK as a substrate. An autoradiograph of the $^{32}$P incorporated into MEK and immunoblot analysis of the BRAF levels are shown, indicating that the kinase activity of the Sf9-produced complexes increases over time during the incubation reaction, whereas the activity of the Tni-FNL-produced complexes remains constant. (*C*) Kinase activity of Tni-FNL-produced BRAF:14-3-3$_2$:MEK complexes incubated alone or with GppNHp-KRAS-FMe (7 molar excess), 30% PS liposomes (41μM), or GppNHp-KRAS-FMe and 30% PS liposomes. The graph represents the average fold increase in kinase activity, with BRAF activity alone set at 1, based on data from 3 independent experiments ± SD. (*B-C*) Also shown are autoradiographs of the $^{32}$P-labeled MEK and immunoblot analyses of BRAF levels from representative experiments.

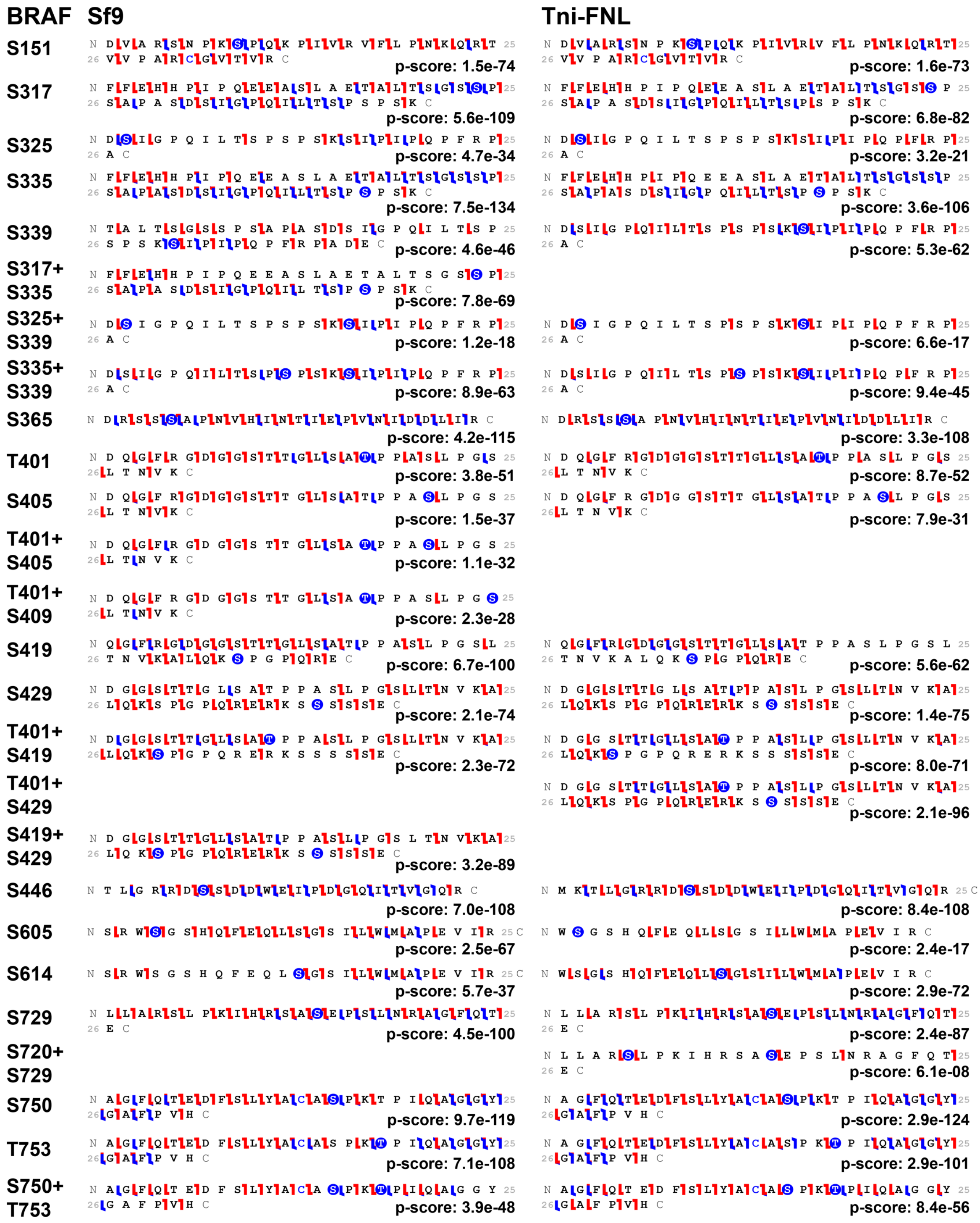

**Fig. S5.** Phosphorylation sites of BRAF in Sf9- and Tni-FNL-produced BRAF:14-3-3$_2$:MEK complexes. Graphical fragment maps are shown, containing the manually verified MS2-based localization for each identified phosphorylation site (blue residues). The maps for Sf9-produced BRAF complexes are on the left, while those for Tni-FNL-produced BRAF complexes are on the right. Flags represent matched fragment ions from electron transfer dissociation (indicated in red) and higher-energy collisional dissociation (indicated in blue). The quality of each identification is indicated by the associated p-score.

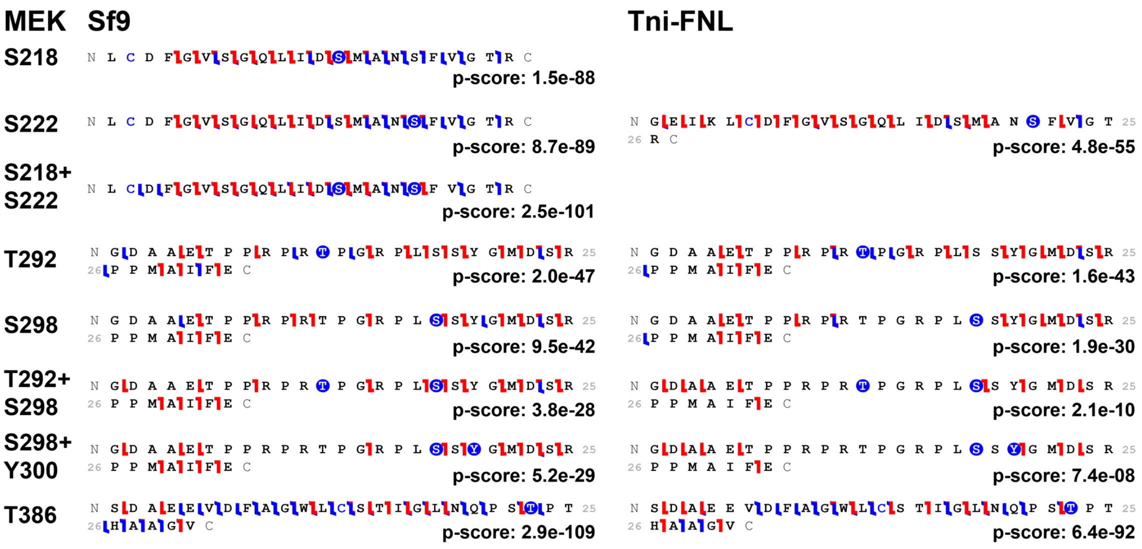

**Fig. S6.** Phosphorylation sites of MEK in Sf9- and Tni-FNL-produced BRAF:14-3-$3_2$:MEK complexes. Graphical fragment maps are shown, containing the manually verified MS2-based localization for each identified phosphorylation site (blue residues). The maps for Sf9-produced MEK are on the left, while those for Tni-FNL-produced MEK are on the right. Flags represent matched fragment ions from electron transfer dissociation (indicated in red) and higher-energy collisional dissociation (indicated in blue). The quality of each identification is indicated by the associated p-score.

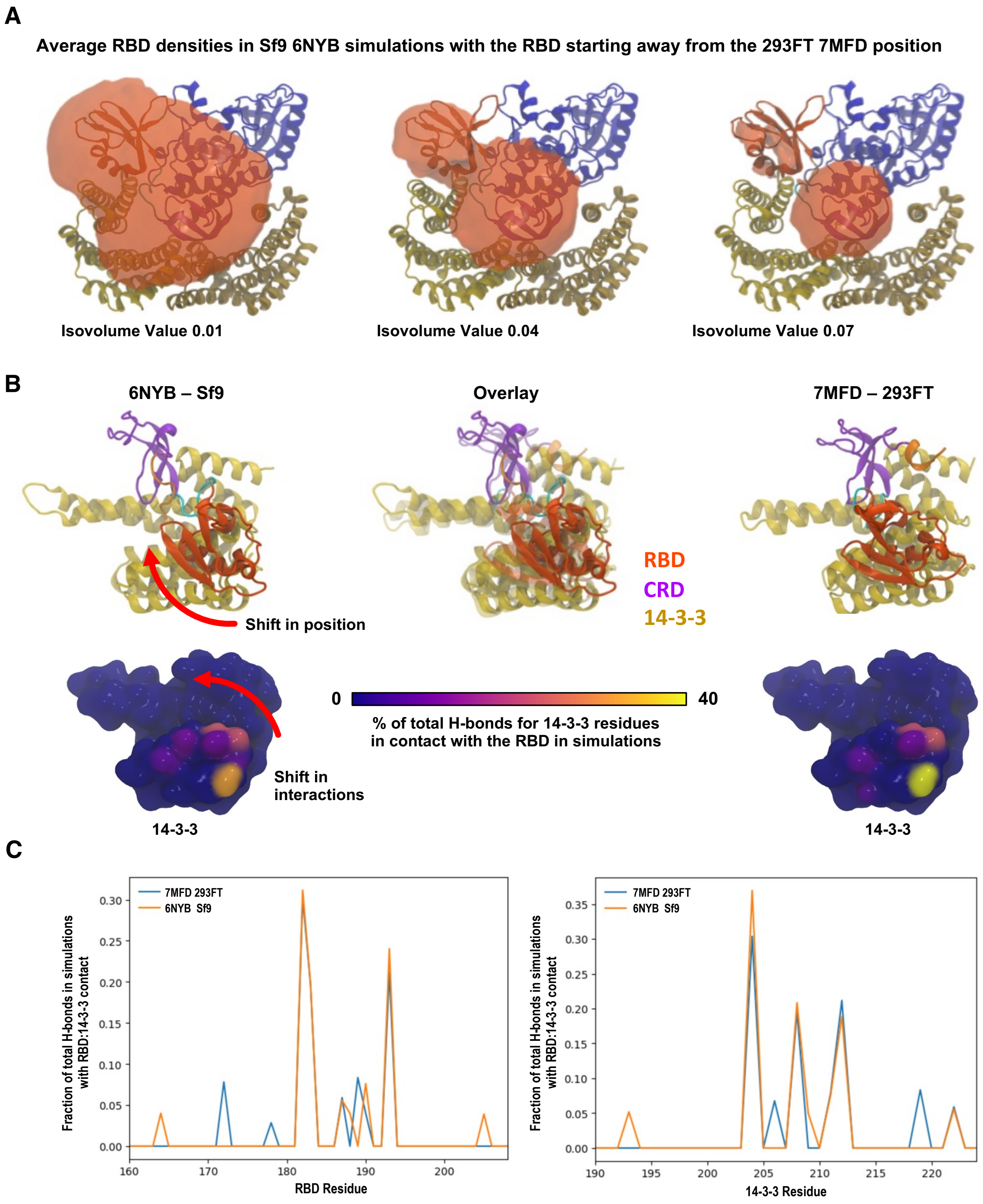

**Fig. S7.** Molecular dynamics simulations of Sf9-produced autoinhibited BRAF complexes. (*A*) To assess the placement RBD in the context of the PDB ID: 6NYB structure, over 600 MD simulations were performed using different starting positions for the RBD and different starting structures for Loop I and II. The volume maps show the average densities of the RBD across all simulations that start with the RBD positioned away from the PDB ID 7MFD location. The volume maps (left-to-right isovolume values of 0.01, 0.04, 0.07) indicate the extent of the large conformational space sampled by the possible RBD configurations as well as several metastable RBD locations, including the conformation observed in 7MFD. (*B*) Spontaneous, unbiased association of the RBD with 14-3-3 is observed in simulations of the SF9 6NYB structures, where the RBD is initially positioned away from the protein. Over the course of the simulation, the RBD settles into a position similar to that seen in the 293FT 7MFD structure (indicated above), with a slight shift that leads to an overlapping, yet distinct, pattern of hydrogen bonding with 14-3-3 (shown below). (*C*) The fraction of total hydrogen bonds formed by each residue in the RBD and the 14-3-3 protomer bound to the pS729 site was calculated based on the aggregated all-atom simulation time for the Sf9 6NYB and 293FT 7MFD structures using only the simulation frames where the RBD is in contact with 14-3-3.

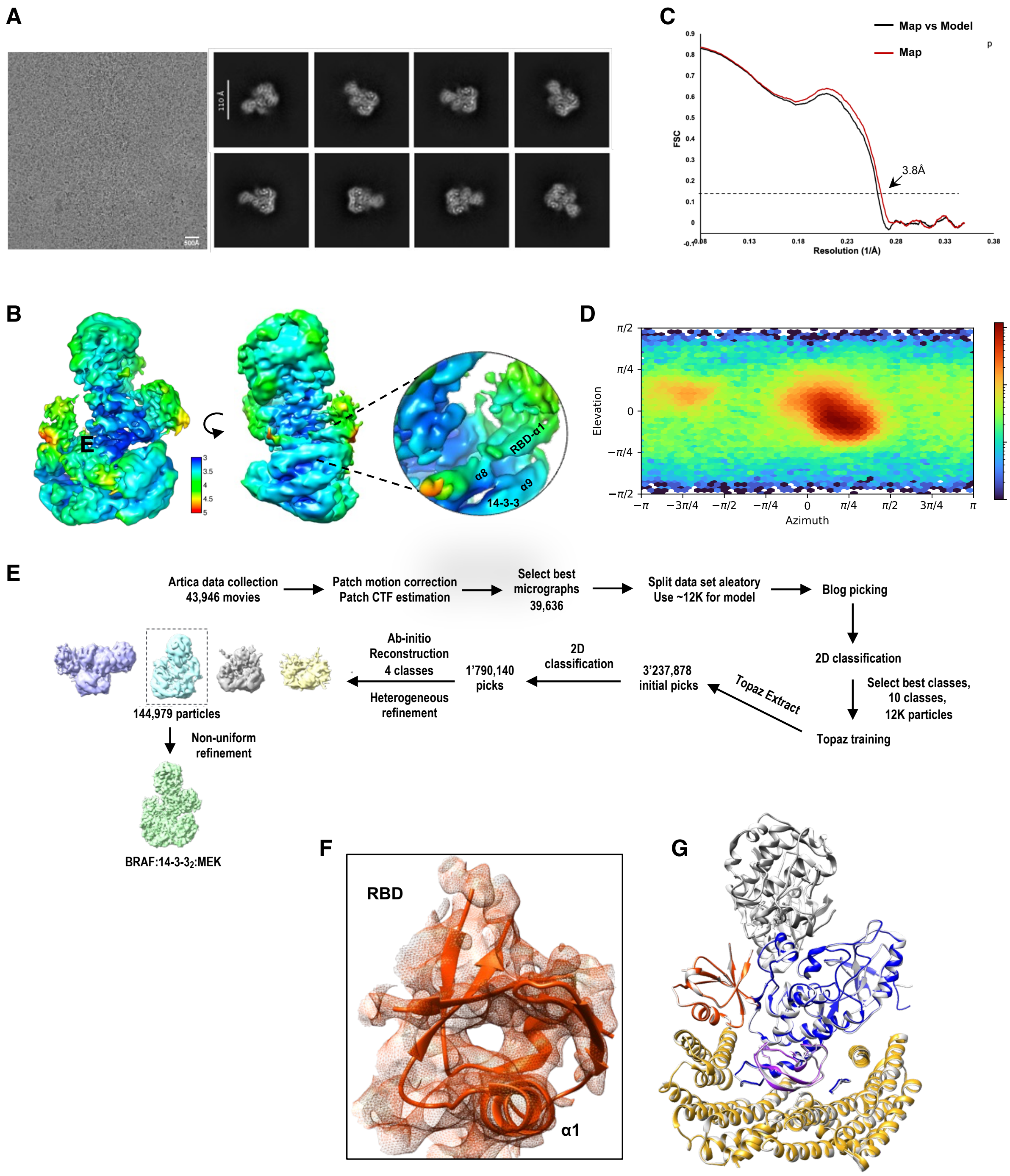

**Fig. S8.** Cryo-EM analysis of Tni-FNL-produced human BRAF:14-3-3$_2$:MEK complexes. (*A*) Representative micrograph and 2D class averages of Tni-FNL-produced BRAF:14-3-3$_2$:MEK particles are shown. (*B*) BRAF:14-3-3$_2$:MEK cryo-EM map colored according to local resolution. (*C*) Map-model FSC curve, verifying the model fit to the cryo-EM density map. (*D*) Angular distribution plot of particle orientations for the global non-uniform refinement. (*E*)

Schematic of the cryo-EM 3D reconstruction workflow. (*F*) Density map showing the fit of the RBD from the Tni-FNL-produced complexes. (*G*) Superimposition of the structures of the BRAF:14-3-$3_2$:MEK complex produced in Tni-FNL insect cells and in 293FT mammalian cells (PBID: 7MFD, gray), with a Cα R.M.S.D. of 0.36 Å.

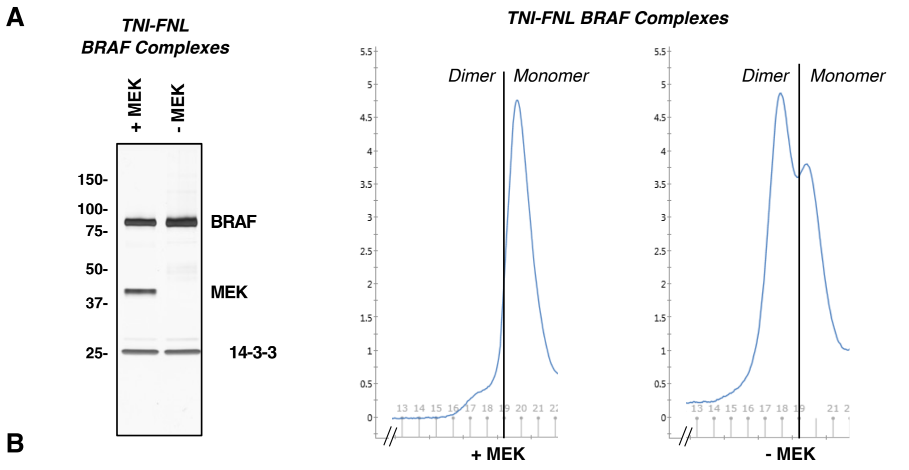

| Residue | Tni-FNL BRAF (Monomer) | Tni-FNL BRAF (Dimer) | Sequence Surrounding the Modifed Reside |
|---|---|---|---|
| S151 | 13.28% | 11.66% | DVARSNPKSPQKPIVRVFLPNKQRTVVPARCGVTVR |
| S317 | 1.29% | 2.39% | FFEHHPIPQEEASLAETALTSGSSPSAPASDSIGPQILTSPSPSK |
| S325 | 0.55% | 1.53% | DSIGPQILTSPSPSKSIPIP QPFRPA |
| S335 | 0.80% | 1.87% | FFEHHPIPQEEASLAETALTSGSSPSAPASDSIGPQILTSPSPSK |
| S339 | 2.45% | 9.63% | TALTSGSSPSAPASDSIGPQILTSPSPSKSIPIPQPFRPADE |
| S365 | 96.59% | 90.95% | SSSAPNVHINTIEPVNIDDLIR/DRSSSAPNVHINTIEPVNIDDLIR |
| T401 | 31.07% | 34.65% | DQGFRGDGGSTTGLSATPPASLPGSLTNVK |
| S405 | 1.40% | 3.36% | DQGFRGDGGSTTGLSATPPASLPGSLTNVK |
| S419 | 6.53% | 5.91% | QGFRGDGGSTTGLSATPPASLPGSLTNVKALQKSPGPQRE |
| S429 | 48.86% | 33.13% | DGGSTTGLSATPPASLPGSL TNVKALQKSPGPQRERKSSSSE |
| S446 | 80.08% | 80.17% | RDSSDDWEIPDGQITVGQR/TLGRRDSSDDWEIPDGQITVGQR/MKTLGRRDSSDDWEIPDGQITVGQR |
| S465 | 0.26% | 0.36% | IGSGSFGTVYK |
| S605 | 1.68% | 3.08% | WSGSHQFEQLSGSILWMAPEVIR/SRWSGSHQFEQLSGSILWMAPEVIR |
| S614 | 1.84% | 5.63% | WSGSHQFEQLSGSILWMAPEVIR/SRWSGSHQFEQLSGSILWMAPEVIR |
| S729 | 98.46% | 99.54% | LLARSLPKIHRSASEPSLNRAGFQTE |
| S750 | 3.31% | 2.96% | AGFQTEDFSLYACASPKTPIQAGGYGAFPVH/AGFQTEDFSLYACASPKTPIQAGGYGAFPVH |
| T753 | 2.18% | 1.45% | AGFQTEDFSLYACASPKTPIQAGGYGAFPVH/AGFQTEDFSLYACASPKTPIQAGGYGAFPVH |

**Fig. S9.** Analysis of Tni-FNL autoinhibited monomer and activated dimer BRAF complexes. (*A*) BRAF complexes co-expressed with 14-3-3 alone or with 14-3-3 and MEK1 were analyzed by SDS-PAGE followed by silver staining. (*B*) Gel filtration analysis showing that BRAF complexes co-expressed with MEK1 elute primarily as monomers (Fraction 20), whereas BRAF complexes not co-expressed with MEK1 elute primarily as dimers (Fraction 18). (*C*) BRAF proteins from Tni-FNL-expressed monomeric and dimeric complexes were evaluated for phosphorylation status using quantitative mass spectrometry. The relative phosphorylation percentage for each detectable site is shown alongside the surrounding sequence. Note that the analysis of monomeric Tni-FNL BRAF shown in Fig. 4C is repeated here, now including all detectable phosphorylation sites, to enable direct comparison with the phosphorylation state of BRAF in activated dimer complexes. Both monomeric and dimeric BRAF proteins were analyzed in the same experiment.

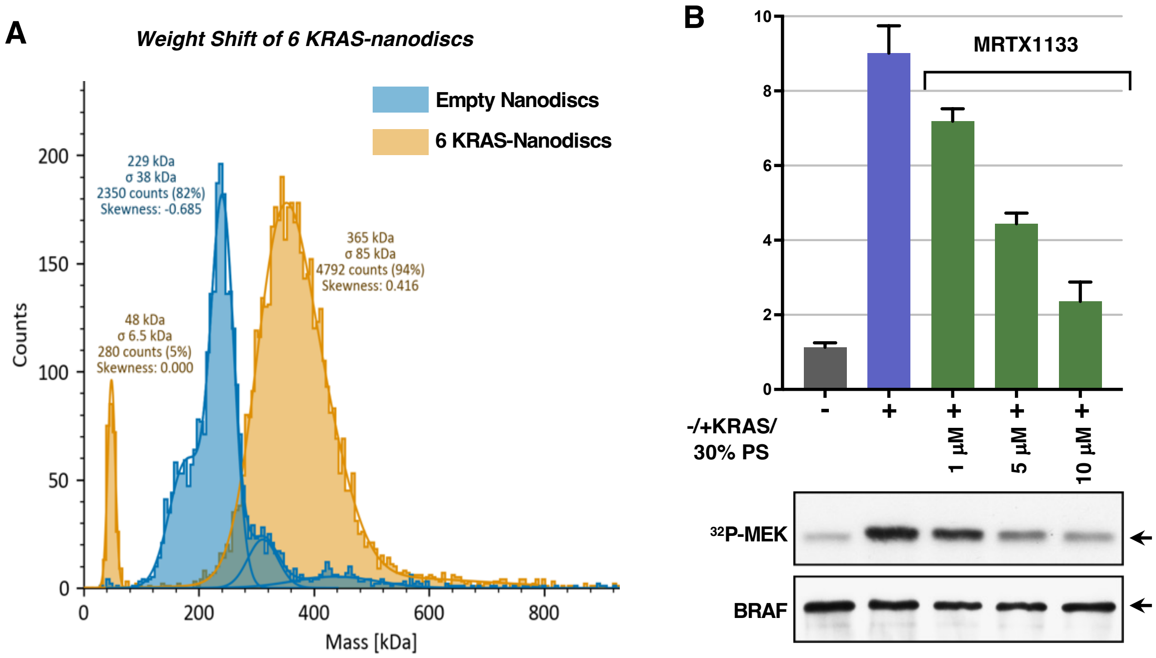

**Fig. S10.** Analysis of KRAS-tethered lipid nanodiscs and the effect of the KRAS inhi bitor MRTX1133 on BRAF activation in vitro. (*A*) Shown is a mass photometry analysis depicting the weight shift after the tethering of KRAS to the lipid nanodiscs. (*B*) Tni-FNL BRAF:14-3-$3_2$:MEK complexes were incubated with GppNHp-KRAS-FMe (7 molar excess) and 30% PS liposomes (83 µM), in the presence or absence of the KRAS inhibitor MRTX1133 as indicated, prior to monitoring for kinase activity in vitro using kinase-dead MEK as a substrate. The graph represents the average fold increase in kinase activity, with BRAF activity alone set at 1, based on data from 3 independent experiments ± SD. Also shown are autoradiographs of the $^{32}$P-labeled MEK and immunoblot analyses of BRAF levels from representative experiments.

**Table S1.** Reagent and resource table

| REAGENT or RESOURCE | SOURCE | IDENTIFIER |
|---|---|---|
| **Antibodies** | | |
| HaloTag mouse monoclonal | Promega | Cat# G9211; RRID:AB_2688011 |
| BRAF F-7 mouse monoclonal | Santa Cruz Biotechnology | Cat# sc-5284; RRID:AB_626760 |
| RAF1 pS259/BRAF pS365 rabbit monoclonal | Abcam | Cat# ab173539; RRID:AB_2813740 |
| pS217/221-MEK rabbit polyclonal | Cell Signaling Technology | Cat# 9121; RRID: AB_331648 |
| MEK 1 mouse monoclonal | BD Biosciences | Cat# 610122; RRID:AB_397528 |
| MEK 2 mouse monoclonal | BD Biosciences | Cat# 610236; RRID:AB_397631 |
| Phospho-MAPK/CDK Substrates (PxS*P or S*PxR/K) rabbit monoclonal | Cell Signaling Technology | Cat# 2325S; RRID:AB_331820 |
| Anti-GST (B-14) | Santa Cruz Biotechnology | sc-138;RRID:AB_627677 |
| **Critical Reagents** | | |
| Superose 6, 10/300 GL | Cytiva | Cat# 29091596 |
| Superdex 200 Increase 10/300 GL | Cytiva | Cat# 28990944 |
| Bio-Beads SM-2 Adsorbents | Bio-Rad Laboratories | Cat# 1523922 |
| Pierce Silver Stain Kit | ThermoFisher Scientific | Cat# 24612 |
| Halolink Resin | Promega | Cat# G1915 |
| Halo-TEV | Promega | Cat# G6602 |
| 1-palmitoyl-2-oleoyl-*sn*-glycero-3-phosphocholine (POPC) | Avanti Polar Lipid | Cat# 850457 |
| 1-palmitoyl-2-oleoyl-*sn*-glycero-3-phospho-L-serine (POPS) | Avanti Polar Lipid | Cat# 840034 |
| 1,2-dioleoyl-sn-glycero-3-phosphoethanolamine-N-[4-(p-maleimidomethyl)cyclohexane-carboxamide] (PEMCC) | Avanti Polar Lipid | Cat# 780201 |
| 1-palmitoyl-2-arachidonoyl-*sn*-glycero-3-phosphocholine (PAPC) | Avanti Polar Lipid | Cat# 850459C |
| 1-palmitoyl-2-oleoyl-*sn*-glycero-3-phosphoethanolamine (PAPE) | Avanti Polar Lipid | Cat# 850757C |
| 1,2-dilinoleoyl-*sn*-glycero-3-phosphoethanolamine (DIPE) | Avanti Polar Lipid | Cat# 850755C |

| | | |
|---|---|---|
| 1-palmitoyl-2-arachidonoyl-*sn*-glycero-3-phospho-L-serine (PAPS) | Avanti Polar Lipid | Cat# 840061C |
| 1-stearoyl-2-arachidonoyl-sn-glycero-3-phospho-(1'-myo-inositol-4',5'-bisphosphate (PI(4,5)P2) | Avanti Polar Lipid | Cat# 850165P |
| N-palmitoyl-D-erythro-sphingo-sylphosphorylcholine (DPSM) | Avanti Polar Lipid | Cat# 850584P |
| Cholesterol | Avanti Polar Lipid | Cat# 700100P |
| $\gamma^{32}$P-ATP | Revvity Health Sciences | Cat# BLU502A001MC |
| **Deposited Data** | | |
| BRAF:14-3-3$_2$:MEK (human) | Martinez Fiesco et al., 2022 (12) | Coordinates: PDBID: 7MFD Cryo-EM map: EMDB: EMD-23813 |
| BRAF:14-3-3$_2$ (human) | Martinez Fiesco et al., 2022 (12) | Coordinates: PDBID: 7MFE Cryo-EM map: EMDB: EMD-23814 |
| BRAF:14-3-3$\zeta_2$:MEK1 (Tni) | This paper | |
| Coordinates of RAS Binding Domain (RBD) of BRAF bound to RAS | Aramini et al., 2015 (29, 30) | PDB ID: 2L05 |
| All simulation parameter files and select simulation files | This paper and Tempkin et al. 2025 (1) | https://bbs.llnl.gov/ |
| **Experimental Models: Cell lines** | | |
| Halo-WT-BRAF 293FT (human) | Martinez Fiesco et al., 2022 (12) | |
| Sf9 (*Spodoptera frugiperda* insect) | ThermoFisher Scientific | Cat# 12659017 RRID:CVCL_0549 |
| Tni-FNL (*Trichoplusia ni*, insect) | Kerafast | Cat# ENH127-FP; RRID:CVCL_RY32 |
| **Recombinant DNA** | | |
| pCMV5-Halo-BRAF$^{WT}$ | NCI-RAS Initiative | N/A |
| **Software and Algorithms** | | |
| EPU Software | ThermoFisher | https://www.thermofisher.com/us/en/home/electron-microscopy/products/software-em-3d-vis/epu-software.html |
| Cryosparc 3.3 | Punjani et al., 2017 (20) | https://cryosparc.com |

| | | |
|---|---|---|
| MotionCor2 | Zheng et al., 2017 (31) | http://msg.ucsf.edu/em/software/motioncor2.html |
| COOT 0.9.6 | Emsley et al., 2010 (25) | https://www2.mrc-lmb.cam.ac.uk/personal/pemsley/coot |
| PHENIX 1.17.1 | Adams et al., 2010 (32) | https://www.phenix-online.org |
| Chimera 1.13.1 | Pettersen et al., 2004 (24) | https://www.cgl.ucsf.edu/chimera |
| Prism 8 | N/A | https://www.graphpad.com/scientific-software/prism |
| Molecular Operating Environment (MOE) | Chemical Computing Group ULC | https://www.chemcomp.com |
| GROMACS | doi 10.1016/j.softx.2015.06.001 | https://www.gromacs.org/ |
| Visual Molecular Dynamics (VMD) | Humphrey et al., 1996 (9) | https://www.ks.uiuc.edu/Research/vmd/ |
| AMBER18 | Salomon-Ferrer et al., 2013 (33) | https://ambermd.org/index.php |
| CHARMM-GUI | Lee et al., 2016 (4) | https://www.charmm-gui.org/ |
| **Other** | | |
| R1.2/1.3 300 mesh Au holey carbon grids | Electron Microscopy Sciences | Cat# Q350AR1.3 |
| Extruder set with heating block | Avanti Polar Lipid | Cat# 610000-1EA |

**Table S2.** Cryo-EM data collection, refinement, and validation statistics

| | **Tni BRAF:14-3-3$_2$:MEK** |
|---|---|
| **Data collection** | |
| Microscope | Talos Arctica |
| Camera | K3 |
| Magnification | 100k, EFTEM mode |
| Voltage (kV) | 200 |
| Electron exposure ($e^-/Å^2$) | 50 |
| Number of frames collected per micrograph | 50 |
| Energy filter slit width | 20eV |
| Defocus range (μm) | -0.8 to -2.5 |
| Pixel size (Å) | 0.81 |
| Movies (no.) | 39,636 |
| Initial particle images (no.) | 3,237,878 |
| Final particle images (no.) | 144,979 |
| Map resolution (Å) | 3.83 |
| FSC threshold | 0.14 |
| Map sharpening B factor ($Å^2$) | 155.8 |
| EMDB code | |
| **Model building and refinement** | |
| Initial model used (PDB) | 7MFD |
| Model composition | |
| Non-hydrogen atoms | 9127 |
| Protein residues | 1147 |
| Protein molecules | 3 |
| Real-space correlation | |
| CCvolume | 0.69 |
| CCmask | 0.70 |
| Mean B factor ($Å^2$) | 75.65 |
| RMS deviations | |
| Bond lengths (Å) (outliers >4σ) | 0.004 (0) |
| Bond angles (°) (outliers >4σ) | 0.820 (14) |
| Validation | |
| MOLProbity score | 2.35 |
| Clashscore | 20.66 |
| Rotamer outliers (%) | 2.0 |
| CaBLAM outliers (%) | 2.3 |
| Cβ outliers (%) | 0.00 |
| Ramachandran plot | |
| Favored (%) | 95.7 |
| Allowed (%) | 4.3 |
| Outliers (%) | 0.05 |
| PDB code | 70053 |

**Dataset S1.** Relative phosphorylation site abundance calculations for phosphorylation at the pS365 and pS729 site for 293FT-produced BRAF proteins used in the in vitro assays.

**Dataset S2.** Relative phosphorylation site abundance calculations for BRAF and MEK produced in Sf9 and Tni-FNL cells.

**Movie S1.** Molecular dynamics simulation of the 293FT BRAF 7MFD structure.
A representative simulation for the 293FT 7FMD structure, showing that the RBD remains in contact with the 14-3-3 protomer bound to the C′ pS729 site. For clarity, MEK, and the unstructured loops were excluded.

**Movie S2.** Molecular dynamics simulation of the Sf9 BRAF 6NYB structure.
A representative simulation for the SF9 6NYB structure, where the RBD is initially positioned away from the protein, demonstrates the spontaneous, unbiased association of the RBD with 14-3-3. For clarity, only the RBD, CRD, the loop connecting the RBD and CRD, and the 14-3-3 protomer bound to the C′ pS729 site are shown.
